# *Mycobacterium tuberculosis* overcomes phosphate starvation by extensively remodelling its lipidome with phosphorus-free lipids

**DOI:** 10.1101/2024.09.27.615480

**Authors:** Robert M. Gray, Debbie M. Hunt, Mariana S. dos Santos, Jiuyu Liu, Aleksandra Agapova, Angela Rodgers, Acely Garza-Garcia, James Macrae, Maximiliano G. Gutierrez, Richard E. Lee, Luiz Pedro S. de Carvalho

## Abstract

Tuberculosis (TB) is the biggest cause of death from infectious disease worldwide [1–4]. The causative agent, *Mycobacterium tuberculosis* (Mtb), possesses a complex cell envelope that determines many of the key physiologic and virulence properties that facilitate infection, which comprises multiple classes of unique lipids [5–7]. The macrophage phagosome is a key reservoir of infection in pulmonary TB and multiple studies have shown that inorganic phosphate (Pi) is limiting in this environment [8–11]. The ability of Mtb to sense and respond to phosphate starvation is required for virulence in animal models and replication in human macrophages in vitro [12–14]. Here, we show that during Pi restriction the Mtb lipidome is markedly remodelled such that phospholipids are replaced with multiple classes of phosphorus-free lipids, some of which have not been documented before. Further, we discover that Mtb can metabolise phospholipid polar heads derived from host pulmonary surfactant as an alternative phosphate source, which sustains cell division for several generations during Pi restriction. These dual manipulations of phospholipid metabolism provide Mtb with an escape from phosphate restriction specific to the infection of alveolar macrophages, one of the earliest steps in establishing pulmonary tuberculosis. The changes in envelope lipidome remodelling, akin to that observed in some marine and terrestrial bacteria [15–20] suggests that standard Mtb culture conditions that use media with high concentrations of Pi do not reflect the physiologic environment during infection, thereby potentially undermining vaccine and drug development for tuberculosis. Moreover, the distinct Mtb phosphate-free lipids and the metabolic pathways that generate them could provide new antibiotic targets.

---

Inorganic phosphate (Pi) is the preferred source of phosphorous for bacteria, an essential component of a plethora of biomolecules. Studies have shown that the Pi starvation response system in Mtb, orchestrated by the two-component system, comprising sensor histidine kinase SenX_3_ and transcriptional response regulator RegX_3_, are required for full virulence in animal models of TB and are essential for Mtb growth within human macrophages in vitro [8, 12–14]. *Mycobacterium marinum* engineered to fluoresce when encountering a Pi concentration of < 10 µM emits fluorescence from 3 days post infection of zebrafish embryos, and a similarly sensitive reporter in *Salmonella enterica* fluoresces during infection of macrophages in culture but not during infection of epithelial cells, suggesting that Pi starvation is specific to the macrophage phagosome [10, 11].

Environmental and pathogenic bacteria, and some yeasts, have been shown to be able to substitute phospholipids for phosphorus-free lipid species when Pi is limiting [15–22], a strategy also common in plants and green algae [23–26]. Mtb contains a remarkably complex lipidome, organised into a multi-layered envelope composed of an inner plasma membrane (PM) and an outer mycomembrane (mycomembrane or MM) (**Extended Data** Figure 1). The enzymatic pathway for the breakdown of phospholipids in Mtb has been proposed to involve the sequential action of yet to be identified phospholipases and a phosphodiesterase, followed by the action of *rv1692*-encoded glycerol-phosphate phosphatase [27]. Mtb is also known to actively transport exogenous phospholipid polar heads across its membrane via the ATP-binding cassette transporter UgpABCE [28, 29], and intriguingly, studies have indicated that this transporter is transcriptionally induced by RegX_3_ [12, 30], suggesting a link between polar head uptake to phosphate starvation. Whilst it is known that Mtb remodels its envelope composition in response to external stimuli such as low oxygen tension or physiologic salinity [31–33], here, we studied the previously uncharacterised enzyme glycerophosphodiesterase 1 (GlpQ1), demonstrating its function in phospholipid remodelling and discovered a dramatic change in envelope lipid composition involving > 1500 lipid species and allowing Mtb to overcome Pi limitation.

## Results

### *glpQ*1 deletion blocks polar head catabolism and disrupts phospholipid remodelling

GlpQ1, encoded by *rv3842*c in the virulent reference strain of Mtb; H37Rv, is annotated by sequence homology to be a glycerophosphodiesterase; an enzyme that hydrolyses the polar lipid-head of glycerophospholipids following their deacylation by phospholipases (**Extended Data** Figure 1b). To probe the enzymatic function of GlpQ1 and its role in PM remodelling we generated an in-frame clean genetic deletion of *glpQ1* in H37Rv (D*glpQ*1 strain) and performed unbiased global metabolomics paired with lipidomics. Parent/wild-type (WT), Δ*glpQ*1 and Δ*glpQ*1*::glpQ*1 strains growing exponentially in liquid culture were subjected to biphasic extraction [34]. The aqueous phase of these extracts contains the cell’s polar metabolites, including the four glycerophosphodiesters of Mtb, the proposed substrates for GlpQ1. Employing a liquid chromatography mass spectrometry (LCMS) protocol hereafter referred to as the HILIC method (hydrophilic interaction liquid chromatography), we quantified 1,064 metabolites in positive ion-mode and 616 in negative ion-mode. Whilst > 96% of features in the Δ*glpQ*1 strain were not significantly altered compared with WT, 3 of the 4 canonical polar-lipid heads of Mtb, glycerophosphoethanolamine (GroPEth), glycerophosphoglycerol (GroPGro) and glycerophosphoinositol (GroPIns) markedly enriched in the Δ*glpQ*1 strain, with fold changes of > 64-fold (2^6^) (**Figure 1a and Extended Data** Figure 2). The 4^th^ lipid head, bisglycerophosphoglycerol (Bis(GroP)Gro), was detected but the chromatographic resolution was poor. We therefore analysed polar cell extracts using a second LCMS method (amide column), which confirmed Bis(GroP)Gro also accumulated markedly in Δ*glpQ*1. (**Figure 1b and Extended Data** Figure 2b). Consistent across both LCMS methods and all four lipid-heads, the phenotype showed partial complementation, with approximately 50 % reversion of lipid head levels to that of the WT in Δ*glpQ*1*::glpQ*1 (**Figure 1b, Extended Data** Figure 2b). These findings strongly support the physiological role of GlpQ1 as a glycerophosphodiesterase of phospholipids in Mtb. By analysing the global lipidome of these strains, we were able to investigate how the marked accumulation of the lipid-heads in the cytosolic compartment translated into PM changes. By using a mycobacterial lipidome platform [6] for analysis of the organic phase of the cell-extracts (**Figure 1c, Extended Data** Figure 3), we detected 3,629 lipids in positive ion-mode, and of these 125 features (3.4 %) were significantly altered in the Δ*glpQ*1 strain. Analysis of the changes seen in the Δ*glpQ*1 strain at the level of phospholipid class (**Figure 1c**) revealed that phosphatidylethanolamine (PE) and cardiolipin (CL) levels were essentially unchanged versus the WT, whereas phosphatidylglycerol (PG) and phosphatidylinositol (PI) levels both fell by approximately 40 % each. Further, these changes showed near total complementation in the Δ*glpQ*1*::glpQ*1 strain.

**Figure 1:**
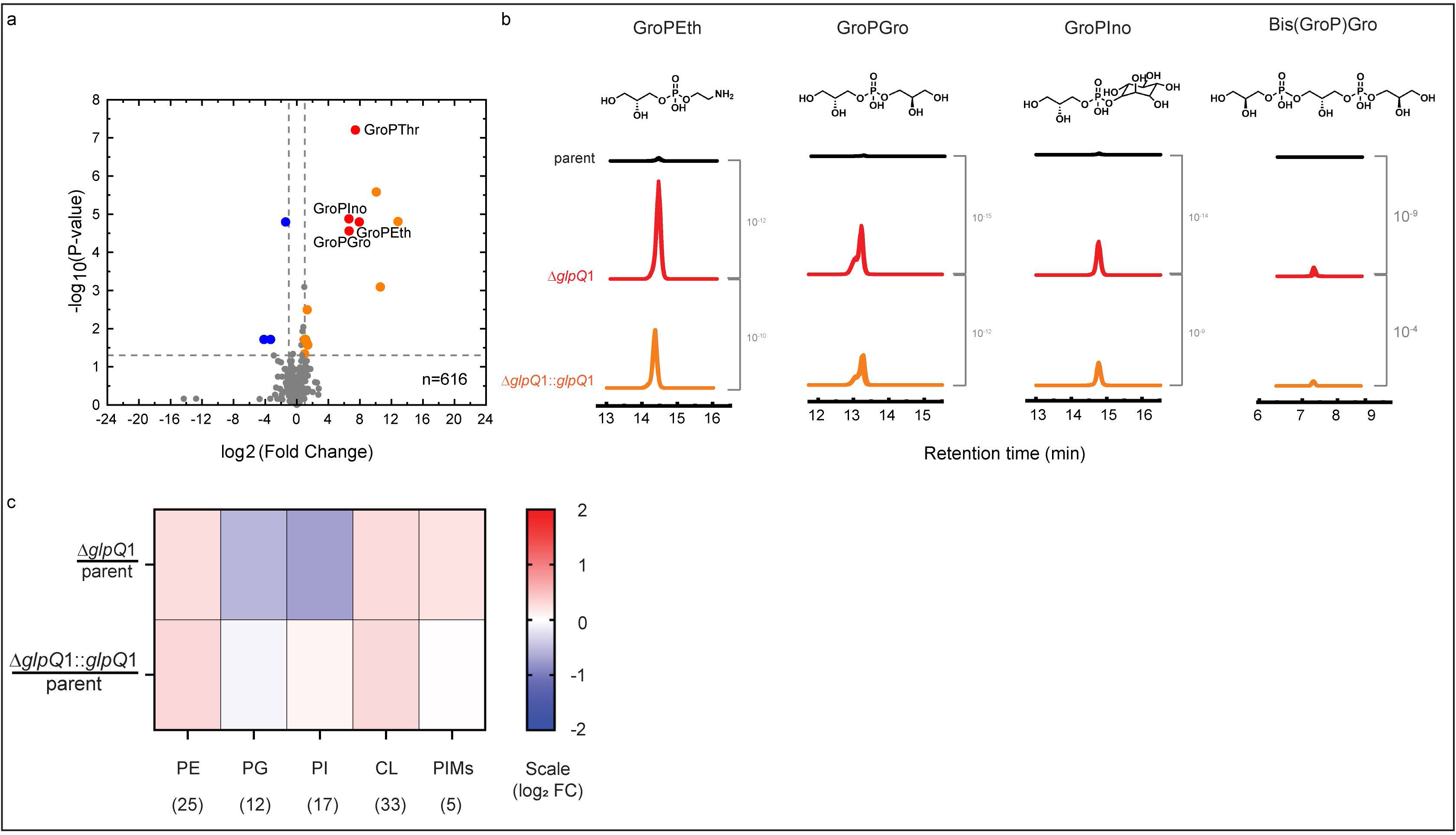
Response of the Mtb metabolome and lipidome to *glpQ*1 deletion. **a:** Volcano plot showing the metabolome of Mtb plotted as the fold change in the mean abundance of each feature in Δ*glpQ*1/parent, showing negative ion-mode. Means were calculated across replicate cultures: Δ*glpQ*1 = 6 replicates, parent = 8 replicates. Features in red are lipid-heads. b: Extracted ion chromatograms for each of the four lipid-heads of Mtb, shown beneath the chemical structure of the neutral molecule. Note that the chromatogram for each replicate culture within each strain are overlain. P values are shown in grey. c: Heat map showing the fold change in the mean abundance of each class of phospholipid in the Δ*glpQ*1 and Δ*glpQ*1::*glpQ*1 strains versus the parent. Blue represents depletion in the mutant, red enrichment. Numbers in brackets represent the number of lipid species in that class.

Measured species of phosphatidylinositol dimannosides (PIMs), a further major component of the Mtb PM, did not alter significantly between the strains (**Figure 1c, Extended Data** Figure 3 **c,d,e).** This is interesting, as PIMs are constructed on a PI membrane anchor, and therefore this conservation of PIM levels in Δ*glpQ*1 suggests that available PI is prioritised for PIM synthesis. Both PI and PIMs are essential in mycobacteria, with loss of viability of *Mycobacterium smegmatis* observed when levels decrease by 70 % and 50 % respectively [35]. The findings that phospholipid levels in Δ*glpQ*1 moderately decrease in 2 of 5 classes and remain constant in 3 of 5, despite all lipid head species markedly accumulating within the metabolome, indicates that robust homeostatic mechanisms must exist at the PM to maintain its composition and function.

### GlpQ1 hydrolyses exogenous lipid heads to maintain growth

Of the 4 remaining features found to be enriched by > 2^6^ in Δ*glpQ*1 in negative-ion mode, collision induced dissociation mass spectrometry (C.I.D.) identified glycerophosphothreonine, a lipid-head not previously observed in Mtb **(Extended Data** Figure 2a**, 2c)**. Phosphatidylthreonine has been described as a minor phospholipid in the PM of mammalian cells [36]. Therefore, GlpQ1 may also metabolise additional lipid-heads, including potentially those from exogenous phospholipids. Consistent with this, in positive ion-mode, we identified glycerophosphocholine (GroPCho) amongst enriched ion-features in the Δ*glpQ*1 strain (**Figure 2a-c).** GroPCho is the lipid-head of phosphatidylcholine (PC), produced after the action of phospholipases, and a major phospholipid of mammalian cells that is also found in some bacteria, but not thought to be present in Mtb. Analysis of the growth media used in the experiments detected multiple PC species, both in the Albumin-Dextrose-Catalase growth supplement, and in the Middlebrook medium itself, reflecting the ubiquitous nature of this phospholipid. Therefore, the accumulation of GroPCho in Δ*glpQ*1 supports Mtb taking up GroPCho derived from PC in the media, which is then hydrolysed by GlpQ1 in the WT (**Figure 2a**).

**Figure 2.**
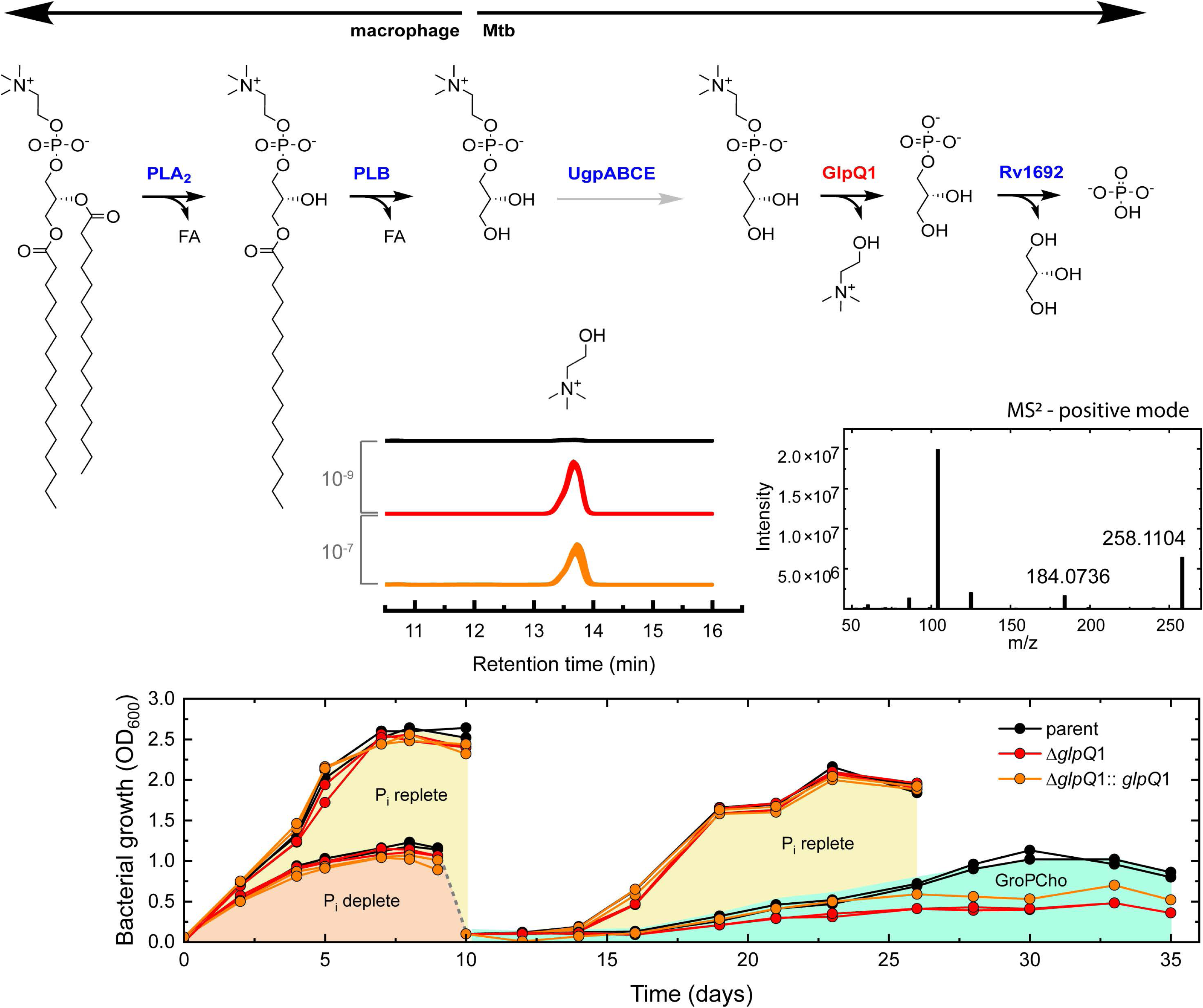
Discovery of glycerophosphocholine as a host-derived alternative phosphate source. **a:** The pathway from host phosphatidylcholine to phosphate, with UgpABCE actively importing the glycerophosphocholine intermediate into the mycobacterium. PLA_2_: phospholipase A2. PLB: phospholipase B. b: Overlay of the EICs for glycerophosphocholine (M+H)^+^ of each replicate culture for each strain, black = WT (6 replicates), red = Δ*glpQ*1 (5 replicates shown), orange = Δ*glpQ*1::*glpQ*1 (5 replicates shown). In the overlays, 1 outlying replicate was removed from the Δ*glpQ*1 and Δ*glpQ*1::*glpQ*1 strains. P values are shown in grey, calculated from all 6 replicate cultures per strain. c: Collision induced dissociation spectrum at the level of MS2 for glycerophosphocholine, (M+H)^+^ in positive ion-mode. The peak at 184 is diagnostic. d. Glycerophosphocholine sole phosphate source experiment. Growth profiles of the strains as labelled in Pi-replete (25 mM) and Pi-free (0 mM) media. At day 9 bacteria were transferred from Pi-free media into fresh Pi-replete media or into GroPCho media (0 mM Pi, 25 mM glycerophosphocholine). Duplicate cultures were performed per strain.

We hypothesised that GlpQ1 degradation of host-derived lipid-heads might provide an alternative source of phosphate to allow Mtb to grow and divide. To test this hypothesis, we transferred exponentially growing bacteria into a modified Pi-free Middlebrook 7H9 medium. Standard Middlebrook medium contains 25 mM Pi, which is highly supraphysiological than the <10 µM Pi present in macrophage phagosomes. After a period of preconditioning, WT, Δ*glpQ*1 and Δ*glpQ1::glpQ*1 strains were grown in Pi-free media. Surprisingly, all three strains could grow in Pi-free media for around eight days, with no substantial difference in growth between the strains (**Figure 2d and Extended Data** Figure 4). This indicates that Mtb must have mobilizable intracellular phosphate stores. In Pi-free culture, growth plateau was reached at a much lower optical density than in the phosphate-replete (25 mM) medium control arm. Next, we transferred bacteria which had reached growth stasis in Pi-free media into fresh Pi-free media supplemented with 25 mM GroPCho. As negative and positive controls, bacteria were also transferred into fresh Pi-free media and into Pi-replete (25 mM) media, respectively. No growth was observed in fresh Pi-free media, whereas exponential growth restarted in 25 mM Pi culture after a lag of 3-4 days, demonstrating that growth stasis was Pi depletion. The bacteria transferred into 25 mM GroPCho, started to divide after a longer lag period of 6 days and at a much slower rate than the bacteria transferred into 25 mM Pi media, reaching a lower plateau at around 20-25 days post transfer. WT Mtb grew better on this substrate than Δ*glpQ*1, with Δ*glpQ*1*::glpQ*1 displaying intermediate growth. These findings demonstrate that Mtb can access exogenous GroPCho as a sole source of phosphate to reactivate cell-division and that this process is partly GlpQ1 dependent.

As GroPCho is the lipid-head of dipalmitoylphosphatidylcholine (DPPC), the main lipid component of human pulmonary surfactant, and that a major site of surfactant breakdown in normal pulmonary physiology is within the phagosome of the alveolar macrophage [37–39], we proposed that GlpQ1 mediated production of phosphate from exogenous GroPCho may be an important nutritional pathway for Mtb during pulmonary infection. No consensus on the importance of *glpQ*1 (*rv3842*c) during infection can be drawn from transposon mutagenesis studies; that is, while DeJesus et al. [40] concluded that disruption of *rv3842*c led to a growth advantage *in vitro*, Sassetti et al. [41] indicated that mutants could survive during murine infection and, in contrast, Rengarajan et al. [8] demonstrated that the gene is required for survival in murine macrophages. We therefore investigated the performance of our clean genetic deletion, Δ*glpQ*1 in the low-dose aerosol infection mouse model of TB **(Extended Data** Figure 5). A 2 log_10_ reduction in pulmonary colony forming units versus the parent strain was observed. Given the partial metabolic complementation observed for of our Δ*glpQ*1::*glpQ*1 strain, we did not include it in this experiment. In a second independent experiment, the phenotype was similar, but of lower magnitude. This experiment suggests that *glpQ*1 is important for Mtb infection.

### Mtb extensively remodels its envelope lipids when grown in the absence of inorganic phosphate

Producing an envelope for daughter cells when dividing in Pi-free conditions poses a substantial phosphate requirement. To overcome this, we hypothesised that Mtb might remodel its lipidome, replacing phospholipids with alternative phosphorus-free lipid classes. To address this, WT, Δ*glpQ*1 and Δ*glpQ*1*::glpQ*1 were cultured in Pi-free medium until growth plateau, then their LCMS lipidome profiles were compared to those of the same strains grown in a 25 mM Pi-replete media. Remarkably, WT Mtb profoundly remodelled its lipidome when cultured in Pi-free medium (**Figure 3a Extended Data** Figure 6). Of 2,963 lipids detected in positive ion-mode, 52 % were significantly altered in the WT when grown in Pi-free vs. in Pi-replete (25 mM) media. Remodelling was roughly symmetrical, with 878 features depleted in Pi-free culture and 674 enriched. Fold changes for the most changed lipids exceeded 2^20^ and the changes were sufficiently extensive as to be visible at the level of the total ion chromatogram **(Extended Data** Figure 6a), with a particularly prominent increase in lipid abundance in the retention time range of 21 and 24 minutes in Pi-free medium-derived lipidome.

**Figure 3.**
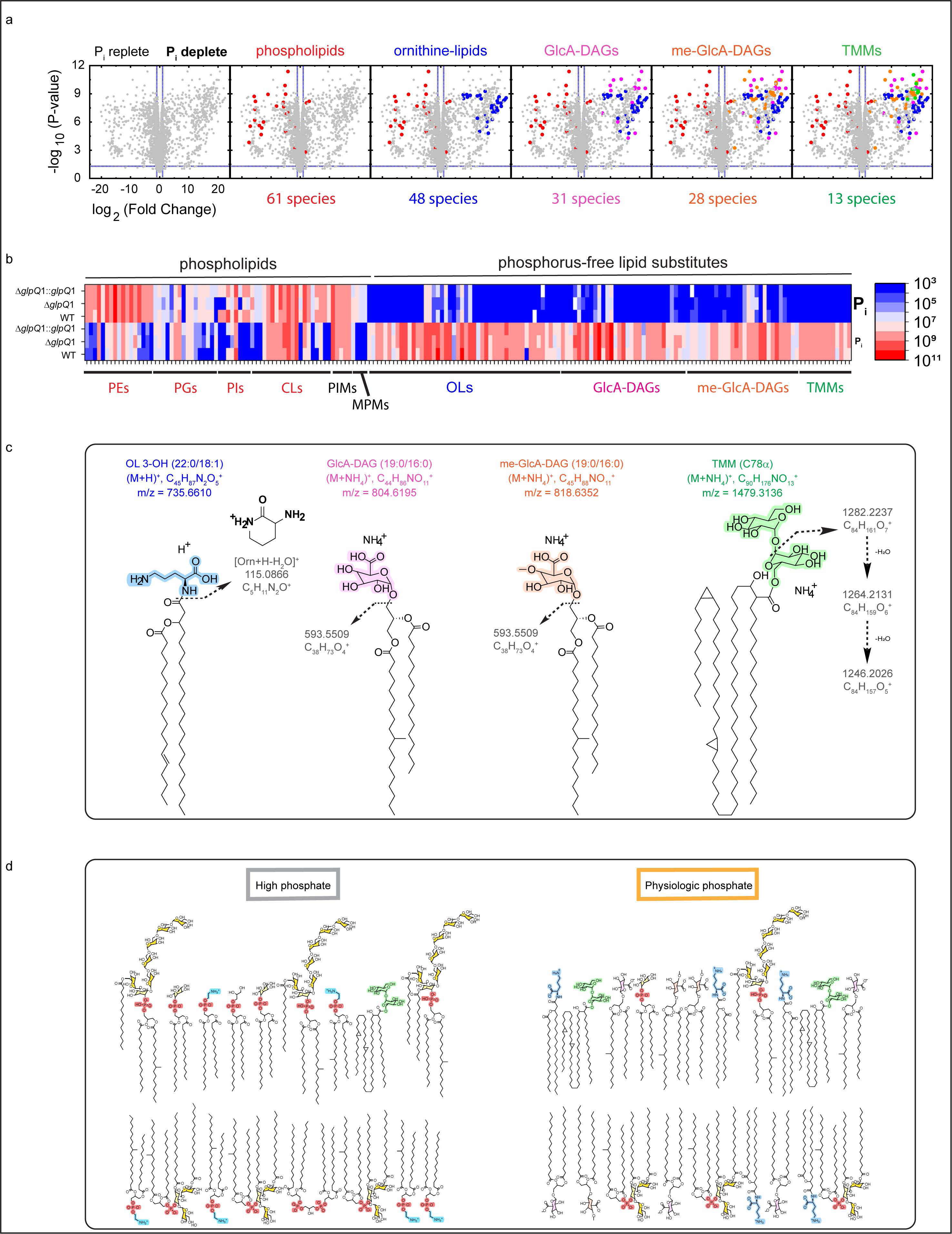
Remodelling of the Mtb lipidome in response to starvation. **a:** Volcano plot showing the lipidome of Mtb, plotted as the fold change in the mean abundance of each detected lipid in the WT grown in Pi-free media/ mean abundance in the WT grown in Pi-replete (25 mM) media, positive ion-mode data. Thus, features to the left of the midline are lipids enriched in replete Pi and features to the right are lipids enriched in zero Pi. Left to right volcanoes stepwise annotate the major classes of altered lipids. Total features plotted = 2963. Means are calculated across 6 replicate cultures (Pi-free culture) and 5 replicates cultures (Pi-replete culture). A CV < 30 in the PBQCs statistical filter has been applied. Representative of 2 independent experiments. b: Heatmap of the abundance of the 181 lipid species comprising the phospholipids and phosphorus-free lipids detected in positive ion-mode, as well as 4 MPMs and 5 PIMs detected in negative ion-mode. Top 3 rows depict the mean abundance of each lipid (area under the curve) in each strain grown in Pi-replete media (25 mM), bottom 3 rows each strain grown in Pi-free media. Lipid species are organised on the x-axis into their classes. Means are calculated across replicate cultures per strain. Representative of two independent experiments. c: Chemical structures for an example of each of the four classes of phosphorus-free replacement lipids. The polar headgroups are coloured, and arrows indicate the major fragmentation lines demonstrated in the C.I.D. data enabling their annotation. d: Diagram of the PM of wild-type Mtb when phosphate is present in excess (left) and when phosphate is restricted (right). Phosphorus-free lipids are drawn with their polar headgroups colour coded as per Figure 3c. Phospholipids are represented as per Extended Data Figure 1b. Mono- and di-acylated PIM_2_ and PIM_6_ are drawn with the mannose sugars coloured in gold and inositol in light yellow, and are labelled in Extended Data Figure 1c.

Furthermore, annotation of significantly altered features based upon accurate mass, retention time and C.I.D. spectra, confirmed the phenomenon of phospholipids being depleted and replaced with phosphorus-free lipid species. These changes were also seen in the Δ*glpQ*1 *and* Δ*glpQ*1*::glpQ*1 strains, indicating that the process is GlpQ independent (**Figure 3b**).

### Substitution of phospholipids by phosphorous-free lipids in Mtb

In Pi-free culture the WT demonstrated a 73 % overall reduction in the abundance of the 61 detected phospholipid species **(Extended Data** Figure 6c), and this figure was remarkably consistent between independent experiments. PIMs, which also contain a phosphate group, were reduced by 33 %. Again, this is a relative sparing of PIM versus PI levels, as PI as a class reduced by 79 %, so suggests shunting of remaining PI to preserve PIMs in the PM. Finally, mannosyl phosphomycoketides (MPMs), a class of mycobacterial phosphate-containing lipids known to be immunogenic and recognized by CD1c-restricted T-cells [42, 43] also proved phosphate sensitive, decreasing by 18-fold in Pi-free culture (**Figure 3b, Extended Data** Figure 6b,c**,e,f)**. Overall, of the 70 phosphate-containing lipid species detected, there was a decrease of abundance of 70% when Mtb was grown in Pi-free medium, demonstrating that Mtb can substantially reduce the phosphate requirement of its cell envelope when challenged by Pi scarcity.

### Identification of the phosphorous-free Mtb lipidome

To maintain cell envelope area and viability, depleted phospholipids must be replaced with alternative, phosphorus-free lipids with similar properties. By using C.I.D. spectral data we were able to identify four classes of phosphorus-free lipids which were markedly enriched in Mtb in response to phosphate starvation, thus acting as substitution lipids.

Ornithine lipids (OLs), have previously been described to replace phospholipids in response to Pi limitation in bacteria, in particular in environmental species [19, 20, 44–48]. OLs containing a b-hydroxy fatty acid in an amide linkage with the α-amino group of ornithine, and the second acyl chain esterified at the β-hydroxy position (**Figure 3c**) are one of the most common OLs, and have been described before in Mtb [49, 50], but when Mtb is cultured in phosphate-replete conditions their abundance is minor and these lipids have seldom been studied in mycobacteria [33], and do not appear in the mycobacterial lipid reference database [6]. In this study, using the presence of a “fingerprint” C.I.D. fragment ion of m/z 115.0866 which corresponds to a 3-amino-2-oxopiperidium ion (**Figure 3c (ii) and** [51]), we confidently annotated 48 species of OL in Mtb grown in Pi-free medium. By contrast, only 10 of these species were detectable in Pi-replete cultured bacteria, and the summed total area under the curve for the OL class showed a 1,020-fold enrichment in the lipidome obtained from Mtb grown in Pi-free medium versus Pi-replete culture Mtb.

Next, we annotated glucuronic acid-containing lipids, glucuronyl-diacylglycerols (GlcA-DAGs). We detected 31 species of GlcA-DAGs as ammoniated ions in our data from the Mtb cultured in Pi-free medium, with only 6 detectable in the Mtb culture in Pi-replete medium. The summed total abundance for GlcA-DAGs showed a mean 4,100-fold increase in lipidomes from Pi-free vs. Pi-replete cultures. C.I.D. confirmed the identity of this class of lipids, with both a neutral loss of 211 seen in positive ion-mode from the ammoniated ions (**Figure 3c**), which represents the loss of the glucuronic acid head group (193) and the ammonium (18) [52], and also a characteristic fragment ion in the MS^2^ spectrum of the deprotonated ion, of mass m/z 249.06, corresponding to a glucuronylglycerol moiety containing an oxirane group [26, 53]. This membrane glycolipid has previously been described in *M. smegmatis* [54], but never studied in Mtb. Again, our data here show that this usually minor, phosphorus-free lipid becomes a dominant lipid class when Mtb is challenged by Pi restriction.

We subsequently found a cluster of 28 ion features that were markedly enriched in lipidomes from Pi-free cultured bacteria and exhibited a retention time of around 4.5 minutes (**Figure 3a, 3b, 3c)**. Examination of their C.I.D. spectra in positive-ion mode indicated that they produced very similar MS^2^ fragmentation to GlcA-DAGs, except the neutral loss seen was now m/z 225 (**Figure 3c**). We hypothesised that as in the case of GlcA-DAGs, this represented the loss of the ammonium adduct and the head group sugar, but rather than glucuronic acid (m/z 193), this moiety now had an m/z 207. This difference of +14 is consistent with a single O-methylation of the glucuronic acid headgroup in these lipids. Therefore, these results represent a novel class of glycolipids, O-methyl glucuronyl-diacylglycerols, or me-GlcA-DAGs.

Methylation of glycans in nature is considered to be rare [55], however, two arguments support this annotation. First, the observed retention time shift in the diol column from 21 minutes for GlcA-DAGs to 4.5 minutes for me-GlcA-DAGs is consistent with the masking of a polar hydroxyl group with a non-polar methyl, which in the diol-column should elute earlier. Second, O-methylated glucuronic acid has been previously documented in several mycobacteria, for instance as a constituent of a pentasaccharide hapten in *M. avium* [56] as well as in glycopeptidolipids in *M. habana* [57]. We therefore consider this annotation as highly plausible.

Assuming our annotations are correct, of 28 putative me-GlcA-DAGs in the Mtb grown in Pi-free medium only 10 were also detectable in the WT grown in Pi-replete media. 22 species had MS^2^ spectra available, all of which showed the neutral loss of 225. The summed total abundance for all 28 me-GlcA-DAGs combined showed a 120-fold increase in Mtb cultured in Pi-free medium vs. in Pi-replete medium. Therefore, me-GlcA-DAGs may be a third phosphorus-free PM lipid which Mtb uses to substitute in for phospholipids.

Surprisingly, we also observed a significant change in trehalose monomycolates (TMMs). These lipids contain long chain mycolic acids, are phosphorus-free, are an essential component of the mycobacterial envelope [58], and have established C.I.D. patterns [6], enabling confident annotation. We detected 13 distinct TMMs as ammoniated ions in Mtb grown in Pi-free medium with each species exhibiting ion intensities of between 10^7^ - 10^8^ counts, whereas none of these lipids were detected in Mtb grown in Pi replete culture (**Figure 3a,b,c)**.

Therefore, during challenge by phosphate restriction, the cell envelope of Mtb undertakes a marked remodelling, with phosphorus-free lipid species which are barely detectable when Pi is freely available becoming dominant classes, married to a reduction in both simple and decorated phospholipids (**Figure 3d).** As such conditions are encountered within the macrophage phagosome, this phosphate restricted lipidome can be considered the physiological makeup.

### A synthetic standard supports the presence of phosphorus-free lipids in Mtb

None of the four phosphorus-free lipid species which we found to be upregulated under Pi depletion is commercially available to use as an LCMS standard, an important experimental validation for novel species. Therefore, we synthesised one of the most abundant species in our Mtb extracts, GlcA-DAG (16:0/16:0) [53] (**Figure 4a, Extended Data** Figure 7). We analysed the synthetic GlcA-DAG (16:0/16:0) lipid standard, which demonstrated comparable negative and positive ion-mode MS^2^ spectral profiles as the same feature in our lipid extracts of Mtb (**Figure 4b, 4c)**. To conclusively prove our annotation of this class of lipid in our Mtb extracts, we analysed an extract from Mtb grown in Pi-free medium, before and after spiking it with synthetic GlcA-DAG (16:0/16:0) to a final concentration of 50 μg/mL. As shown in **Figure 4d**, the extracted ion chromatogram (EIC) for GlcA-DAG 16:0/16:0 (of m/z 762.5726) augmented appropriately in the spiked extract, demonstrating co-elution and therefore confirming the annotation.

**Figure 4.**
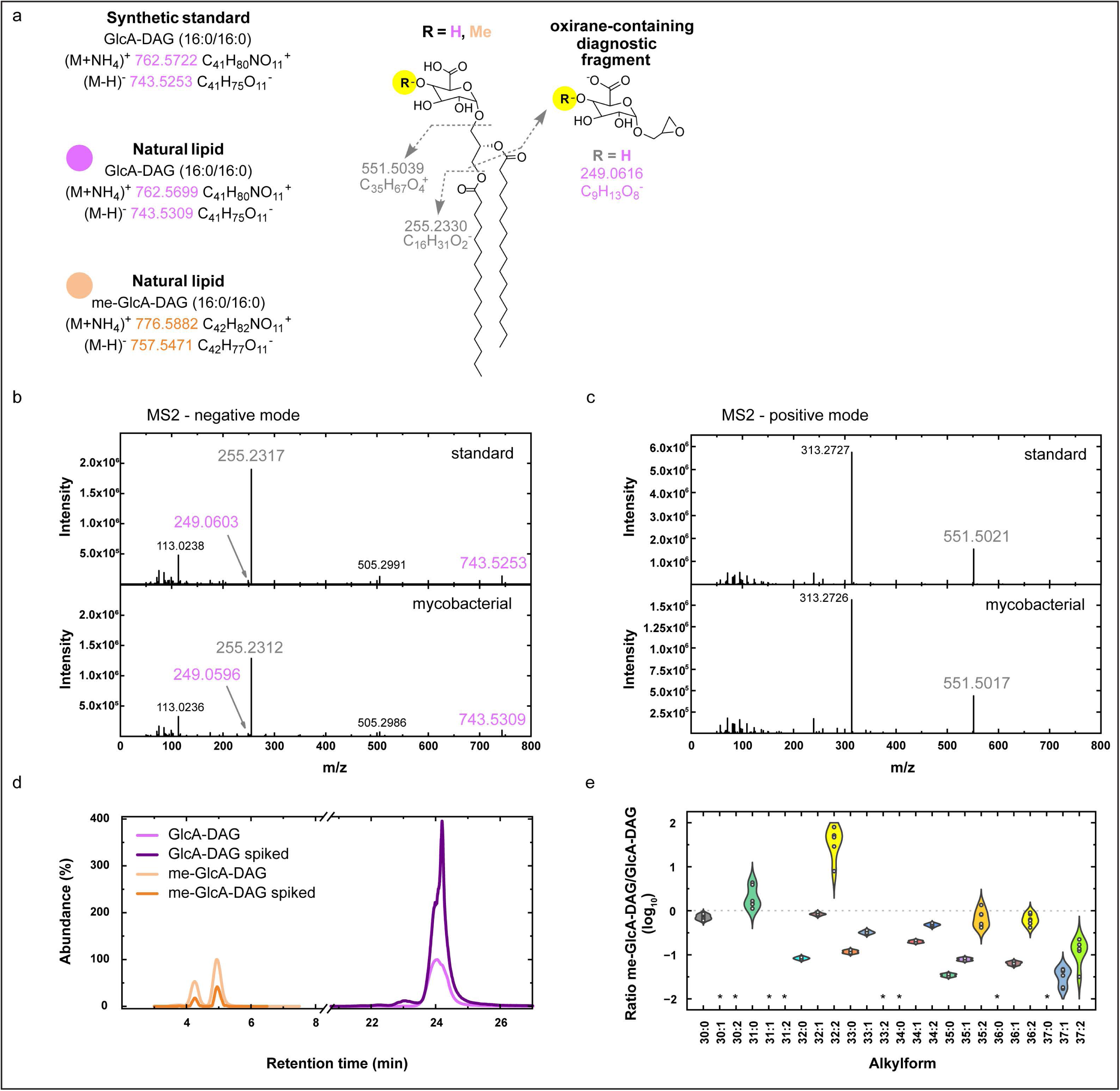
Use of synthetic standard to confirm annotation of glucuronyl-diacylglycerols in Mtb. **a:** Structure of the 16:0/16:0 alkylforms of GlcA-DAG and me-GlcA-DAG, demonstrating the C.I.D. product ions shared by the two lipids (grey) as well as the oxirane-containing diagnostic fragment produced in negative ion-mode MS2 for GlcA-DAGs. The methylation is shown at the 4 OH position of the glucuronic acid sugar, but the position is not confirmed. b: Negative ion-mode C.I.D. MS2 spectra for the deprotonated ion of GlcA-DAG (16:0/16:0) comparing that for the synthetic lipid (top) to that obtained from the mycobacterial lipid extract (bottom). The diagnostic oxirane-containing fragment of m/z 249 is indicated. c: Positive ion-mode MS2 spectra for the ammoniated ion of the same. Note that due to its complete fragmentation at this collision energy, the precursor ion of 762.5726 is not visible. d: Positive ion-mode EICs for the ammoniated ions of me-GlcA-DAG (16:0/16:0) and GlcA-DAG (16:0/16:0) from WT Mtb grown in Pi-free media, before and after spiking with synthetic GlcA-DAG (16:0/16:0) to 50 μg/mL. The synthetic lipid had been stored in 2:1 (v/v) chloroform: methanol for 48 hours prior to spiking. The bifid shape of the peak for the me-GlcA-DAG may indicate that isotypes of this lipid exist with the methylation in 2 different positions upon the sugar, altering the retention time slightly. e: Plot showing the ratio of the abundance of the methylated vs non-methylated glucuronyl-diacylglycerol for each diacylglycerol alkylform detected in Mtb lipid extracts from the WT grown in Pi-free media. The ratio is expressed on a log_10_ scale. Individual points represent the ratio for individual replicates. Asterisks indicate alkylforms where one or both lipids were not detected.

We next leveraged the GlcA-DAG (16:0/16:0) synthetic standard to provide further evidence for the me-GlcA-DAG class in Mtb. It is recognised that extraction from biological samples using methanol-containing solvent mixtures may lead to spurious methylation [59]. We therefore stored the synthetic GlcA-DAG (16:0/16:0) standard in the solvent mixture used in our lipid extraction protocol, 2:1 (v/v) chloroform:methanol, for 48 hours, and then analysed it with LCMS. We could not detect a peak corresponding to a me-GlcA DAG (16:0/16:0) in this sample. Furthermore, spiking of our Mtb cell extract with synthetic GlcA-DAG (16:0/16:0) in 2:1 (v/v) chloroform: methanol to 50 μg/mL did not lead to augmentation of the EIC for the ammoniated ion of me-GlcA DAG (16:0/16:0) (**Figure 4d).** It is therefore unlikely that the proposed me-GlcA-DAG species detected in our Mtb lipidomic experiments are artefactual due to methanol extraction. Further support of this conclusion is provided by the finding that for each diacylglycerol alkylform, the ratio of abundance of the proposed me-GlcA-DAG to the corresponding GlcA-DAG in our Mtb extracts is non constant (**Figure 4e).** Thus, an *in vivo* enzymatic methylation generates these lipids.

## Discussion

Growing bacteria outside of their natural environment poses a fundamental challenge, a metabolic disconnect with reality. In the case of Mtb, the use of media such as Middlebrook 7H9 broth and 7H10/11 agar are rich in glycerol, which is an important carbon source for the bacteria *in vitro*, but absent *in vivo* [60], and Pi is present in supraphysiologic amounts, serving as the pH buffer. Our findings reveal that Pi availability has a profound effect on the composition of mycobacterial lipidomes and membranes.

We characterized extensive lipid remodelling in response to Pi starvation, which revealed two mechanisms by which Mtb overcomes this limitation to cell division. First, Mtb can access host produced phospholipid polar heads, in particular GroPCho, as a source of phosphate, in a GlpQ1-dependent manner. Second, Mtb can extensively replace its PM phospholipids with phosphorus-free lipid species. Both mechanisms are likely important in macrophage infection (where Pi concentrations are below 10 µM) during the pathogenesis of pulmonary TB.

Based on our findings, we propose a model whereby Mtb resides in the phagosomes of alveolar macrophages during human pulmonary infection, which have limited Pi but are rich in GroPCho. GroPCho is produced from DPPC which is also taken up by phagosomes as part of normal pulmonary surfactant metabolism [37–39]. Either host or secreted Mtb phospholipases may hydrolyse DPPC to GroPCho. After active uptake into Mtb by the ABC transporter UgpABCE, GlpQ1 hydrolyses host derived GroPCho to provide an alternative source of phosphate. The Mtb genome encodes only 4 ATP powered transporters specific for carbohydrates in its PM, in marked contrast to *M. smegmatis*, which contains 19. [61, 62]. This is postulated to be a consequence of Mtb encountering the carbohydrate poor environment of the phagosome during infection. That Mtb contains the UgpABCE transporter specific for glycerophosphodiesters therefore suggests this is an important source of nutrition during such macrophage infection. Interestingly, nuclear magnetic resonance studies have shown PC levels decrease and GroPCho levels increase over time inside ex vivo lung granulomas from Mtb infected guinea pigs [63].

During Pi restriction Mtb can generate ATP from cytosolic polyphosphate stores and this mechanism may be required for the full virulence of Mtb in guinea pigs and for survival in THP-1 macrophages [64]. In parallel, we show here that Mtb reduces the amounts of PE, PI, CL, PIMs and MPMs in its membranes. In this way, the cell envelope may be considered a second accessible phosphate storage compartment for Mtb, resulting in phospholipids being replaced by phosphorus-free lipids. These lipids are upregulated to maintain a viable PM, but the observed increase in TMMs, which shuttle mycolic acids into the outer mycomembrane [5], suggests that remodelling in response to low Pi extends beyond the PM.

Our findings are important for understanding the composition of the Mtb cell envelope under physiologic conditions that reflect Pi concentrations in alveolar macrophages. We show that > 50 % of lipids in the Mtb lipidome are significantly altered during culture in Pi-free media, with 2^20^ fold increases in individual lipids. This previously unrecognized aspect of the Pi starvation response involving phospholipid substitution could allow Mtb survival during the early phases of infection and may modulate immunogenicity of the pathogen. Thus, culturing Mtb in Pi-replete media, which is the current standard, may result in key differences, particularly in experiments aimed at developing vaccines and antibiotics against TB. This has important consequences for current efforts to produce lipid-based vaccines against Mtb [65–69], as lipids known to date to be presented by CD1 molecules to T-cells may not necessarily be of high abundance in the Pi-starved lipidome. MPMs are a direct example of this. Conversely, immunologists and vaccinologists have not studied the immune response to physiologic, phosphate-poor Mtb. These experiments are likely to yield important differences, as we already know that ornithine-containing lipids and glucuronyl-containing lipids are immune modulatory [70–73]. Additionally, identifying the biosynthetic enzymes responsible for assembling phosphorous-free lipids could reveal targets for drug development.

## Online Methods Strains

To generate the Δ*glpQ1* knock-out strain from the parent H37Rv, an unmarked, in-frame deletion of the *rv3842c* gene was made using the method of Parish and Stoker [74]. Briefly, 1.5 kb of flanking sequence of the genes *rv3842c* (*glpQ1*) was amplified from the genomic DNA of H37Rv with KAPA HiFI Hotstart taq using the primers Rv3842c 5’ for and Rv3842c 5’ rev for the 5’ side and Rv3842c 3’ for and Rv3842c 3’rev for the 3’ side (see table below). The products were cloned into pCR4blunt (Invitrogen) and sequenced. The inserts were digested with HindIII and XbaI and the fragments ligated together using T4 DNA ligase. The ligation product was amplified using primers Rv3842c 5’ for and Rv3842c 3’ rev and the fragment was cut with HindIII and cloned into the HindIII site of p2NIL. The PacI fragment of pGOAL 17 containing the *lacZ* and *sacB* genes was cloned into the PacI site of the resulting plasmid to make the Δ*glpQ1* construct. Competent H37Rv cells were prepared from cells grown to an OD_600_ of 1 and washed with 10 % glycerol and electroporated with 2 μL plasmid. Single crossovers were selected on 7H11 plates containing kanamycin and Xgal. The blue colonies were then streaked on 7H11 plates containing sucrose and Xgal, and the resulting white colonies were screened for double crossovers. Successful gene deletion was confirmed by next generation sequencing using the Francis Crick Institute’s Advanced Sequencing Science Technology Platform.

The construct for complementation (pML1335-Pimyc-rv3842c) was generated by isothermal assembly (NEBuilder HiFi) using the plasmid pML1335, a gift from Michael Niederweis (Addgene plasmid # 32377 [75]) Firstly, we replaced the promoter of pML1335 with the intermediate promoter Pimyc (https://pubmed.ncbi.nlm.nih.gov/15687379/). Secondly, we replaced the *gfpm2+* insert with the *rv3842c* gene amplified from Mtb H37Rv genomic DNA. Competent Δ*glpQ1* cells were electroporated with pBS-Int (for integrase) and with pML1335-Pimyc-rv3842c, and then transferred to 5 mL of 7H9 medium with ADC growth supplement and tyloxapol, and left standing overnight at 37 °C. The electroporated cells were plated on 7H11 medium with OADC growth supplement containing 100 mg/mL hygromycin B to select for complemented colonies. Colonies were screened for successful complementation construct insertion by PCR with DNA obtained by InstaGene Matrix (BioRad)

**Table 1.**
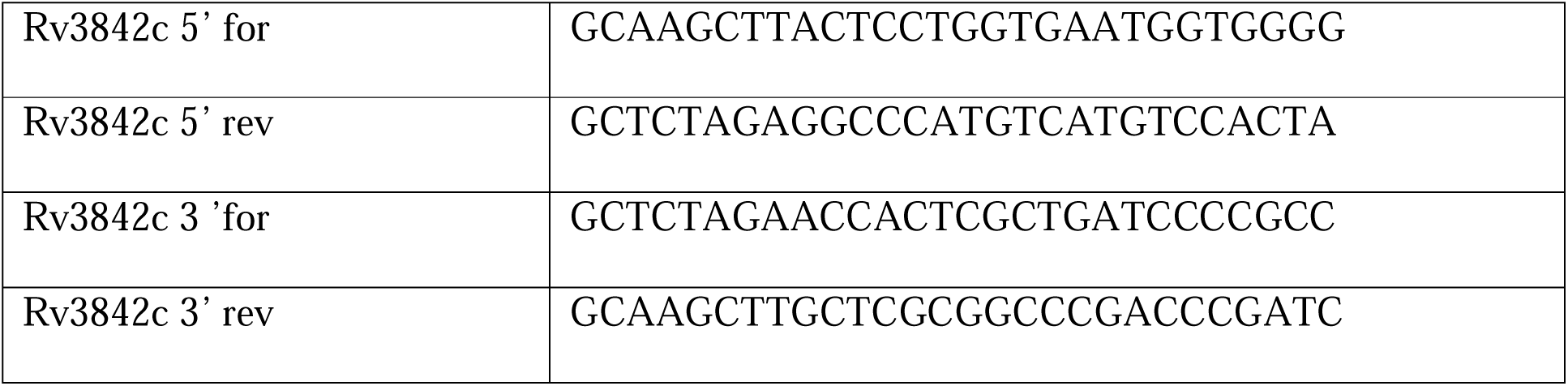
Primers used for *rv3842c* Gene Deletion.

### LC-MS Methods

All solvents used were Optima® HPLC grade from Sigma Aldrich.

#### HILIC Method: ZIC-pHILIC column Chromatography – Orbitrap MS

Cultures were grown in Middlebrook 7H9 complete media until OD_600_ = 1 and then 1 mL transferred onto 0.22 μm filter paper and grown for 5 days on Middlebrook 7H10 media with ADC supplement. After 5 days, cells were scraped from filters into 2:1 (v/v) chloroform: methanol and placed on dry ice for 1 hour to inactive the Mtb. Samples were then removed from the containment level 3 laboratory. Samples were then incubated at 4 °C for one hour with regular sonication, centrifuged at 16,000 rpm, then the supernatant transferred into an Eppendorf tube and dried under nitrogen stream. The pellet was re-extracted with 450 µL methanol: water (2:1, (v/v), containing 3 nmol 13C5, 15N1-valine for LC-MS), and added to the extract. The dried extract was then suspended a final time in 50 µL chloroform, 150 µL methanol, and 150 µL water. Once settled, the upper phase of this biphasic mixture consisted the aqueous sample containing the polar metabolites.

The LC-MS method used was adapted from [76] and is as per published previously [77]. Samples were injected into a Dionex UltiMate LC system (Thermo Scientific) with a ZIC-pHILIC column (Merck Sequant). A 15-min elution gradient of 20% solvent B was used, and this was followed by a 5-min wash of 95:5 solvent A to solvent B and 5-min re-equilibration; solvent B was acetonitrile and solvent A was 20 mM ammonium carbonate in water. Other parameters were as follows: flow rate 300 µL/min; column temperature 25°C; injection volume 10 µL; autosampler temperature 4°C. MS was performed with positive/negative polarity switching using an Q Exactive Orbitrap (Thermo Scientific) with a HESI II (Heated electrospray ionization) probe. MS parameters were as follows: spray voltage 3.5 kV and 3.2 kV for positive and negative modes, respectively; probe temperature 320°C; sheath and auxiliary gases were 30 and 5 arbitrary units, respectively; full scan range: 70 to 1050 m/z with settings of auto gain control (AGC) target and resolution as Balanced and High (3 x 10^6^ and 70,000), respectively. Data were recorded using Xcalibur 3.0.63 software (Thermo Scientific). Mass calibration was performed for both ESI polarities before analysis using the standard Thermo Scientific Calmix solution. To enhance calibration stability, lock-mass correction was also applied to each analytical run using ubiquitous low-mass contaminants. Parallel reaction monitoring (PRM) acquisition parameters: resolution 17,500, auto gain control target 2 × 10^5^, maximum isolation time 100 ms, isolation window m/z 0.4; collision energies were set individually in HCD (high-energy collisional dissociation) mode. PBQC samples were prepared by pooling equal volumes of each sample and were analysed throughout the run to provide a measurement of the stability of the system. To confirm the identification of significant features, PBQCs were analysed in ddMS2 mode. Data were acquired using Xcalibur 3.0.63 (Thermo Fisher Scientific) and Progenesis (Nonlinear Dynamics) was used for data alignment and peak detection. Data from samples were normalized against the total ion abundance. Annotations were assigned to accurate masses with a maximum error of 5 ppm using the Chemspider database and confirmed with MS2 spectra where available. Freestyle 1.7 SP2 (Thermo Scientific) was used to manually check the integration of all the peaks of the glycerophosphodiesterase species and to review MS2 data.

#### Amide Method: Amide column Chromatography – Time of Flight MS

Cultures were grown in Middlebrook 7H9 complete media until OD_600_ = 1, and then 1ml transferred onto 0.22 μm filter paper and grown for 5 days on Middlebrook 7H10 media with ADC supplement. After 5 days, cells were scraped from filters into a 1ml volume of 2:2:1 solution (v/v/v) of acetonitrile: methanol: water with glass beads, and lysed via bead beating with MP FastPrep 24 at 6.5 m/s for 30 secs, then filtered using Corning Spin-X centrifuge tubes, then transferred to LC-MS vials. LCLMS analysis was performed on an Agilent 1290 LC system coupled to an Agilent G6230B ToF mass spectrometer. Chromatography was performed using a Waters XBridge Amide column, 4.6 x 100 mm maintained at 40 °C. Solvent A was 95 % 20 mM ammonium hydroxide, 20 mM ammonium acetate, pH 9.0; 5 % acetonitrile. Solvent B was acetonitrile. Analytes were eluted using a flow rate of 0.4 mL/min and the following mobile phase gradient: 0 – 3 min, 85 – 30 % B; 3 – 12 min, 30 –2 % B; 12 – 15 min, 2 % B; 15 – 16 min, 2 – 85 % B. The injection volume was 20 μL. The Agilent G6230B ToF was operated in positive and negative polarities with ESI using a dual AJS ESI source. Capillary, nozzle and fragmentor voltages were set at 3500 V, 2000 V and 110 V respectively. Data were collected in the m/z range 50 – 1200 and saved in centroid mode. Dynamic mass axis calibration was achieved by continuous infusion of a reference mass solution, which enabled accurate mass spectral measurements with an error of less than 5 ppm. Data from samples were normalized against the total ion abundance. Peak detection was performed with Agilent MassHunter.

#### Diol Method: BETASIL diol column Chromatography-Orbitrap MS

For the lipidomic analyses of exponentially growing bacteria in standard Pi culture conditions, Mtb strains were grown in 4 mL of detergent-free Middlebrook 7H9 complete media until OD_600_ = 1. For zero phosphate lipidomics, Mtb strains were preconditioned with 72-hours of growth in Pi-free media or 25 mM Pi media, then transferred into fresh media of the same Pi concentration, detergent free, 24 mL volume in 50 mL Falcon^TM^ tubes, and grown until growth stasis-defined as 48-hours with static OD_600_ measurements. The composition of the modified Middlebrook media is as details in the *Sole Phosphate Source Experiments* section below.

For extractions, in both experiments, 4 mL of cell culture was then spun for 10 minutes at 3,000 rpm and supernatant discarded. For the zero phosphate experiments, at this stage a proportion of the cell pellet from the cultures grown in 25 mM phosphate was discarded in order to normalise cell mass between the experiment’s two arms. The pellets were then washed with Optima® water twice, then extracted in 2:1 (v/v) chloroform: methanol, and incubated on dry ice for one hour to inactivate the Mtb. Samples were removed from the containment level 3 laboratory and then incubated at 4 °C with regular sonication for one hour then spun at 16,000 rpm for ten minutes, and supernatant transferred and dried under nitrogen stream. The pellets were re-extracted with 1500 μL 2:1 (v/v) methanol: chloroform, sonicated, centrifuged at 16,000 rpm and added to the dried extracts. Combined lipids were dried under nitrogen and the dried pellet dissolved in 100 μL of 1:1 (v/v) chloroform: methanol. This cell lipid extract was diluted 1:2 with solvent A (70:30 (v/v), hexanes: isopropanol, 0.02% (m/v) formic acid, 0.01% (m/v) ammonium hydroxide), centrifuged at 1,500 rpm for 5 minutes to remove trace non-lipidic materials, then transferred to a glass autosampler vial (Agilent). The LC–MS method was adapted from [6]. Samples were injected onto a BETASIL diol column (5 μm x 150 mm x 2.1 mm, with BETASIL diol guard column (10 mm x 2.1 mm), held at 20°C) in an Ultimate 3000 HPLC system coupled to a Thermo Q Exactive Orbitrap MS. Lipids were eluted at 0.15 mL/min with a binary gradient from 0 % to 100 % solvent B (70:30 (v/v) isopropanol: methanol, 0.02 % (m/v) formic acid, 0.01 % (m/v) ammonium hydroxide): 0–10 min, 0 % B; 17–22 min, 50 % B; 30–35 min, 100 % B; 40–44 min, 0 % B, followed by additional 6 min 0 % B post-run. MS data were acquired in both polarities using a full scan method. The positive and negative HESI-II spray voltages were 4.5 and 3.5 kV, respectively; the heated capillary temperature was 250 °C; the sheath gas pressure was 30 psi; the auxiliary gas setting was 20 psi; and the heated vaporizer temperature was 150 °C. The parameters of the full mass scan were as follows: a resolution of 70,000, an auto-gain control target under 3 x 10^6^, a maximum isolation time of 200 ms, and an m/z range 200–3000. To confirm the identification of significant features, PBQCs were ran in data-dependent top-N (ddMS2-topN) mode; parameters as follows: a resolution of 17,500, an auto gain control target under 2 x 10^5^, a maximum isolation time of 100 ms, an isolation window of m/z 0.4 and normalized collision energy of 35 V. Data were acquired using Xcalibur 3.0.63 (Thermo Fisher Scientific). Progenesis (Nonlinear Dynamics) was used for data alignment and peak detection. Sample data were normalized against the total ion abundance. Annotations were assigned to accurate masses with a maximum error of 10 ppm using the MycoMass database [6], and confirmed with MS2 where available and as described in the text. Freestyle 1.7 SP2 (Thermo Scientific) was used for further analysis.

#### Analysis of synthetic GlcA-DAG (16:0/16:0) standard

The Diol Method was run with total cell lipid extract from WT Mtb grown in Pi-free modified media. One aliquot of this extract was ran unspiked, and one aliquot after spiking the extract to a final concentration of 50 μg/ mL with synthetic GlcA-DAG (16:0/16:0) which had been stored first for 48-hours in 2:1 (v/v) chloroform: methanol. The two samples were run in the diol-orbitrap LC-MS system, and a gain in the abundance of the ion feature corresponding to the ammoniated ion of GlcA-DAG (16:0/16:0) was observed in positive-ion mode.

Additionally, the total cell extract and the synthetic standard were analysed side-by-side under the same PRM conditions: resolution 17,500, auto gain control target 2 × 10^5^, maximum isolation time 100 ms, isolation window m/z 0.4; collision energies were set in HCD mode at HCDs of 10.00, 20.00 and 30.00. Retention times and MS2 spectra were a match in both positive and negative-ion modes.

### Synthesis of GlcA DAG (16:0/16:0)

Synthesis was guided by [53]. All solvents used for chromatography were purchased from Fisher Scientific. Flash column chromatography silica cartridges were obtained from Biotage Inc. The purification of mixtures was conducted using Biotage Isolera 1. ^1^H NMR spectra were recorded on a Varian INOVA-500 spectrometer or on a Bruker 400 MHz NMR spectrometer. Chemical shifts (δ) are reported in parts per million (ppm) relative to the residual solvent peak, while coupling constants (J) are reported in hertz (Hz). Waters ACQUITY UPLC H Class Bio coupled with Xevo G2-S QToF with UNIFI was used for High resolution Mass analysis (HRMS). Refer to Extended Data Figure 7 for the synthesis scheme.

### (S)-4,4’-(((3-(allyloxy)propane-1,2-diyl)bis(oxy))bis(methylene))bis(methoxybenzene) (2)

To a solution of (R)-3-(allyloxy)propane-1,2-diol (**1,** 1.0 g, 7.57 mmol) in DMF (20 mL) was added sodium hydride (470.0 mg, 95 %, 19.67 mmol) at 0°C. After stirring for 15 min, p-methoxybenzyl chloride (3.08 g, 19.67 mmol) and a catalytic amount of tetra-n-butylammonium iodide (TBAI) were added, and the mixture was allowed to warm to room temperature. After stirring overnight at 60 °C, the reaction mixture was quenched with MeOH, and then ethyl acetate was added. The organic solution was washed with water, saturated NaHCO_3_ and brine, then dried over MgSO_4_, filtered, and concentrated. The residue was applied onto a column of silica gel. Elution with ethyl acetate/hexane and subsequent concentration of the appropriate fractions gave **2** as an oil (2.0 g, 5.37 mmol, 71 %). MS (ESI): m/z 373 (M^+^+H). ^1^H NMR (400 MHz, CDCl_3_) δ 7.33 – 7.22 (m, 4H), 6.94 – 6.82 (m, 4H), 5.89 (ddt, *J* = 17.3, 10.5, 5.5 Hz, 1H), 5.26 (dq, *J* = 17.3, 1.7 Hz, 1H), 5.21 – 5.12 (m, 1H), 4.62 (s, 2H), 4.47 (s, 2H), 4.01 – 3.98 (m, 2H), 3.81 (s, 3H), 3.80 (s, 3H), 3.75 (dt, *J* = 5.6, 4.7 Hz, 1H), 3.65 – 3.47 (m, 4H).

### (S)-2,3-bis((4-methoxybenzyl)oxy)propan-1-ol (3)

A mixture of **2** (370.0 mg, 0.99 mmol), PdCl_2_ (17.6 mg, 0.99 mmol), and CuCl (98.0 mg, 0.99 mmol) in DMF/water (10/1, v/v, 11 mL) was stirred overnight under nitrogen. The mixture was then diluted with diethyl ether (10 mL) and filtered over Celite. The filtrate was washed with water (5 mL), and the layers were separated. The organic phase was dried over MgSO_4_, filtered, and concentrated in vacuo. The residual oil was purified by silica gel column chromatography (eluent: ethyl acetate/hexanes) to afford **3** (144.0 mg, 0.43 mmol, 44 %); MS (ESI): m/z 333 (M^+^+H). ^1^H NMR (400 MHz, CDCl_3_) δ 7.27 – 7.23 (m, 4H), 6.94 – 6.81 (m, 4H), 4.66 – 4.51 (m, 2H), 4.47 (d, *J* = 2.1 Hz, 2H), 3.81 (s, 3H), 3.80 (s, 3H), 3.75 – 3.53 (m, 5H).

### ((2R,3R,4S,5R,6S)-3,4,5-tris(benzyloxy)-6-((S)-2,3-bis((4-methoxybenzyl)oxy)propoxy) tetrahydro-2H-pyran-2-yl)methyl acetate (5)

Freshly made **4**((2R,3R,4S,5R,6R)-3,4,5-tris(benzyloxy)-6-iodotetrahydro-2H-pyran-2-yl)methyl acetate, 1.53 g, 2.55 mmol)[53] was dissolved in dry CH_2_Cl_2_ (25 mL) and cannulated into a stirred mixture of **3** (470.0 mg, 1.41 mmol), tetrabutylammonium iodide (TBAI, 1.83 g, 4.96 mmol), 2,4,6-tri-tert-butylpyrimidine (969.0 mg, 3.90 mmol), and freshly activated 4 Å molecular sieves in dry CH_2_Cl_2_ (20 mL). The reaction was stirred for 4 days at room temperature under N_2_. The mixture was diluted with Ethyl acetate (20 ml), quenched with 1 M Na_2_S_2_O_3_, and then filtered through Celite. The filtrate was washed with water and brine, dried over MgSO_4_, and concentrated under reduced pressure. The residue was diluted with Et_2_O (20 ml), and the remaining TBAI was excluded by filtration. The filtrate was concentrated, and the residue was purified by flash chromatography (eluent: ethyl acetate/hexanes) to give **5** (687.0 mg, 0.85 mmol, 60%). MS (ESI): m/z 807 (M^+^+H). ^1^H NMR (400 MHz, CDCl_3_) δ 7.36 – 7.13 (m, 19H), 6.86 – 6.70 (m, 4H), 4.96 (d, *J* = 10.8 Hz, 1H), 4.86 – 4.73 (m, 3H), 4.70 – 4.49 (m, 5H), 4.44 – 4.39 (m, 2H), 4.20 – 4.09 (m, 2H), 3.96 (dd, *J* = 9.6, 8.9 Hz, 1H), 3.82 – 3.76 (m, 1H), 3.76 – 3.73 (m, 1H), 3.72 (s, 3H), 3.71 (s, 3H), 3.57 – 3.40 (m, 5H), 1.94 (s, 3H).

### ((2R,3R,4S,5R,6S)-3,4,5-tris(benzyloxy)-6-((S)-2,3-bis((4-methoxybenzyl)oxy)propoxy) tetrahydro-2H-pyran-2-yl)methanol (6)

NaOMe in MeOH (25% wt., 350 μL) was added to a stirred solution of **5** (400.0 mg, 0.49 mmol) in CH_2_Cl_2_ (5 mL). The mixture was stirred for half an hour and then neutralized with Amberlite 120R (H^+^ form) resin. The mixture was filtered, eluent concentrated, and the residue was purified by flash chromatography (eluent: ethyl acetate/hexanes) to obtain **6** (350.0 mg, 0.46 mmol, 92%). MS (ESI): m/z 787 (M^+^+Na).^1^H NMR (400 MHz, CDCl_3_) δ 7.39 – 7.18 (m, 20H), 6.91 – 6.72 (m, 4H), 4.98 (d, *J* = 10.9 Hz, 1H), 4.93 – 4.78 (m, 3H), 4.77 – 4.57 (m, 5H), 4.45 (s, 2H), 3.99 (t, *J* = 9.3 Hz, 1H), 3.80 (dt, *J* = 5.6, 2.6 Hz, 2H), 3.78 (s, 3H), 3.77 (s, 3H), 3.70 – 3.64 (m, 2H), 3.59 – 3.48 (m, 5H).

### (2S,3S,4S,5R,6S)-3,4,5-tris(benzyloxy)-6-((S)-2,3-bis((4-methoxybenzyl)oxy)propoxy) tetrahydro-2H-pyran-2-carboxylic acid (7)

**6** (950.0 mg, 1.24 mmol), 2,2,6,6-tetramethyl-1-(l1-oxidaneyl)piperidine (TEMPO, 100.0 mg, 0.64 mmol), and (Diacetoxyiodo)benzene (BAIB, 2068.0 mg, 6.42 mmol) in a mixture of CH_2_Cl_2_/H_2_O (2:1, 12 mL) was stirred at room temperature for 3 h. The reaction was quenched with 0.25 M Na_2_S_2_O_3,_ and the resulting biphase was extracted with ethyl acetate (15 ml). The organic layer was washed with water, dried over MgSO_4_, and concentrated under reduced pressure. The residue was purified by flash chromatography (eluent: ethyl acetate/hexanes) to obtain **7** (850.0 mg, 1.09 mmol, 88%). MS (ESI): m/z 777 (M^+^-H) ^1^H NMR (400 MHz, CDCl_3_) δ 7.39 – 7.16 (m, 19H), 6.87 – 6.79 (m, 4H), 4.97 (d, *J* = 10.9 Hz, 1H), 4.88 – 4.79 (m, 3H), 4.74 – 4.66 (m, 2H), 4.65 – 4.58 (m, 3H), 4.46 (s, 2H), 4.28 (d, *J* = 10.1 Hz, 1H), 4.05 – 3.94 (m, 1H), 3.88 – 3.78 (m, 2H), 3.77 (s, 3H), 3.76 (s, 3H), 3.70 (dd, *J* = 10.2, 8.8 Hz, 1H), 3.62 – 3.54 (m, 4H).

### benzyl (2S,3S,4S,5R,6S)-3,4,5-tris(benzyloxy)-6-((S)-2,3-bis((4-methoxybenzyl)oxy) propoxy)tetrahydro-2H-pyran-2-carboxylate (8)

Benzyl alcohol (138.0 mg, 1.28 mmol) was added to a stirred mixture of **7** (663.0 mg, 0.85 mmol), 2-(1H-benzotriazol-1-yl)-1,1,3,3-tetramethyluronium hexafluorophosphate (HBTU, 646.0 mg, 1.70 mmol), N,N-Diisopropylethylamine (DIPEA, 275.0 mg, 2.13 mmol), and N,N-dimethylpyridin-4-amine (DMAP, 41.6 mg, 0.34 mmol) in CH_2_Cl_2_ (10 mL). The mixture was stirred overnight at room temperature and then quenched with water. The organic layer was then concentrated under reduced pressure, and the residue was purified by flash chromatography (eluent: ethyl acetate/hexanes) to obtain **8** (250.0 mg, 0.29 mmol, 34%). MS (ESI): m/z 892 (M^+^+Na). ^1^H NMR (400 MHz, CDCl_3_) δ 7.44 – 7.07 (m, 24H), 6.87 – 6.75 (m, 4H), 5.14 (s, 2H), 4.97 – 4.84 (m, 2H), 4.82 – 4.68 (m, 3H), 4.65 – 4.55 (m, 3H), 4.44 (d, *J* = 9.1 Hz, 3H), 4.36 – 4.27 (m, 1H), 3.98 (t, *J* = 9.3 Hz, 1H), 3.87 – 3.77 (m, 3H), 3.76 (s, 3H), 3.75 (s, 3H), 3.75 – 3.70 (m, 1H), 3.62 – 3.52 (m, 3H), 2.17 (s, 5H).

### benzyl (2S,3S,4S,5R,6S)-3,4,5-tris(benzyloxy)-6-((S)-2,3-dihydroxypropoxy)tetrahydro-2H-pyran-2-carboxylate (9)

Ceric ammonium nitrate (CAN, 1794.0 mg, 3.27 mmol) was added to a stirred solution of **8** (948.0 mg, 1.09 mmol) in MeCN/H_2_O (11:1, 12 mL) at room temperature. The mixture was stirred at room temperature for 2 h, then diluted with ethyl acetate (20 ml) and H_2_O to create a biphase. The organic layer was washed with water and brine, dried over MgSO_4_, and concentrated under reduced pressure. The residue to obtain crude **9** (300.0 mg, 0.48 mmol, 44%). MS (ESI): m/z 629 (M^+^+H). ^1^H NMR (400 MHz, Chloroform-*d*) δ 7.30 – 7.26 (m, 5H), 7.26 – 7.22 (m, 6H), 7.22 – 7.18 (m, 7H), 7.09 – 7.04 (m, 2H), 5.16 – 5.04 (m, 2H), 4.85 (d, *J* = 10.92 Hz, 1H), 4.79 – 4.71 (m, 2H), 4.71 – 4.61 (m, 2H), 4.57 (d, *J* = 11.80 Hz, 1H), 4.39 (d, *J* = 10.74 Hz, 1H), 4.23 (d, *J* = 9.92 Hz, 1H), 3.90 (t, *J* = 9.35 Hz, 1H), 3.87 – 3.74 (m, 2H), 3.72 – 3.64 (m, 2H), 3.54 (dt, *J* = 9.62, 5.04 Hz, 2H), 3.41 – 3.32 (m, 1H).

### (2S,3S,4S,5R,6S)-6-((R)-2,3-bis(palmitoyloxy)propoxy)-3,4,5-trihydroxytetrahydro-2H-pyran-2-carboxylic acid (GlcA DAG 16:0/16:0, (10))

Palmitic acid (53.0 mg, 0.207 mmol), 4-(6-cyano-2-methyl-7-oxo-4,8-dioxa-2,5-diazadec-5-en-3-ylidene)morpholin-4-ium hexafluorophosphate(V) (COMU, 89.0 mg, 0.21 mmol), and DMAP (38.0 mg, 0.31 mmol) was added to a stirred solution of crude **9** (65.0 mg, 0.10 mmol) in DMF (5 mL) and the mixture stirred under N_2_ for 24 h. A second equal portion of palmitic acid, COMU, and DMAP was then added as above, and the mixture was stirred for a further 24 h to drive the reaction to completion. The mixture was then diluted with ethyl acetate (10 ml) and washed with saturated NaHCO_3_, water, and brine. Then dried over MgSO4 and concentrated under reduced pressure. The residue was purified by flash chromatography (eluent: ethyl acetate/hexanes) to obtain the intermediate ester, which was taken directly forward to deprotection. A mixture of Pd(OH)_2_/C (20%, 10.0 mg), intermediate ester (10.0 mg, 9.05 μmol) in MeOH/THF (3:2, 5 mL) containing acetic acid (40 μL) was stirred under a H_2_ atmosphere for 2 h. Then degassed and filtered through a layer of Celite and the filtrate was concentrated under reduced pressure. Final purification by silica gel chromatography of the residue (eluent: CHCl_3_/MeOH) gave the product **10** (GlcA DAG 16:0/16:0, 6.0 mg, 8.05 μmol, 8%). ^1^H NMR (500 MHz, DMSO-*d*_6_) δ 5.11 (td, *J* = 8.17, 5.31 Hz, 1H), 4.94 (d, *J* = 5.11 Hz, 1H), 4.83 (d, *J* = 6.55 Hz, 1H), 4.70 (t, *J* = 2.89 Hz, 1H), 4.38 – 4.27 (m, 1H), 4.20 – 4.13 (m, 1H), 3.86 – 3.75 (m, 1H), 3.72 – 3.66 (m, 1H), 3.56 – 3.51 (m, 1H), 3.38 (q, *J* = 7.04 Hz, 2H), 3.23 (ddd, *J* = 9.71, 6.35, 3.72 Hz, 1H), 2.27 (td, *J* = 7.31, 2.96 Hz, 4H), 1.23 (s, 52H), 0.85 (t, *J* = 6.82 Hz, 6H). HRMS-ESI calculated for C_41_H_75_O_11_ [M-H]^-^ 743.53094, found: 743.53131.

### Glycerophosphocholine Sole Phosphate Source Experiments

Pi-free and 25 mM Pi-replete modified media were constituted based upon Middlebrook 7H9 media. The base for both modified media had the following composition:

Per 900 mL base:

Ammonium sulphate 0.5 g L-Glutamic Acid 0.5 g Sodium citrate 0.1 g

Ferric ammonium citrate 0.04 g MgSO_4_ 0.05 g

KCl 0.547 g

0.1M pyridoxine 59 µL 0.1M biotin 20.8 µL 0.1M CaCl_2_ 45 µL

0.1M ZnSO_4_ 34.7 µL

0.1M CuSO_4_ 62.5 µL

1:2 glycerol (50%) 4 mL MOPS buffer 22 g

dH_2_O 896 mL

From these 900 mL aliquots, to prepare Pi-free media, 1.46 g NaCl was added. To make Pi-replete (25 mM) media, 3.0 g of NaH_2_PO_4_ was instead added.

Tyloxapol was added to each to a final concentration of 0.05 %, and pH was adjusted to pH 6.6 using concentrated NaOH. Finally, 100 mL of 10x ADC growth supplement was added to make up a final volume of 1 L. Media were filter sterilised through a 0.22 μm membrane.

Bacteria were grown in 7H9(c) until OD_600_ = 1. These were used to inoculate pre-conditioning cultures, of 24 mL volume in Falcon^TM^ conical centrifuge tubes, at a starting OD_600_ = 0.06. For the Pi-free arm this stage was Pi-free medium, for the high-Pi arm this was Pi-replete (25 mM) medium. These were incubated at 37 °C with continuous rotation for 72-hours. Bacteria were then transferred into 100 mL of fresh media of the same Pi concentration as in the pre-culture stage, at a starting OD_600_ = 0.06, in roller bottles, and growth curves commenced.

Once the Pi-free cultures had reached growth stasis, defined as 48-hours at a static OD_600,_ bacteria from these cultures were transferred into fresh Pi-free medium (negative control), fresh Pi-replete (25 mM) medium (positive control) or into fresh Pi-free medium to which GroPCho had been added to a final concentration of 25 mM. These cultures were all of 24 mL volume in 50 mL Falcon^TM^ tubes. These were incubated at 37 °C with continuous rotation. Each condition/strain was performed in duplicate cultures and the experiment was performed twice.

### Mouse Infection Experiments

All infection studies were approved by the Francis Crick Institute Ethics Committee and performed under a UK Home Office approved Animal License (P4D8F6075) by Angela Roberts. C57BL/6 mice were bred and maintained in the Biological Research Facility at the Francis Crick Institute. Procedures involving mice were performed in strict accordance with the United Kingdom Animals (Scientific Procedures) Act 1986 and the Institute’s policies on the Care, Welfare and Treatment of Animals.

Female C57BL/6 mice at 6-8 weeks old were utilised for all of the infection studies. Mice were exposed to low-dose aerosolised TB infection in an infection chamber using a modified Glas-Col nebuliser system (Glas-Col, Terre Haute, USA). WT H37Rv and deletion strains were grown to mid log-phase in Middlebrook 7H9 media containing ADC and then diluted to produce an infection dose of approximately 100 CFU/mouse lung. Immediately after infection lungs were removed from 5 mice/infection group to assess the infection load CFU. Thereafter mycobacterial CFU are assessed in the lungs of 5 mice/infection group at set time-points after infection: 30, 60, 90 and 120 days.

For CFU analysis, whole lungs were homogenised in tubes containing a solution of saline/0.01% digitonin (Merck, CAS number 11024-24-1) with sterile 3 mm glass beads (Merck) using a FastPrep-24 homogenisation system (MP-Biomedicals Inc.). Lung homogenates were serially diluted 10-fold, and duplicate samples plated onto Middlebrook 7H11 agar plates containing OADC supplement. Plates were incubated at 37 °C and assessed from day 14 post plating.

### PDIM extraction and Thin Layer Chromatography (TLC)

20 mL of bacterial culture for each strain (Middlebrook 7H9(c) media at OD_600_ = 0.2) was centrifuged at 3000 rpm for 5 minutes. Supernatant was discarded and the pellet resuspended in 1 mL of phosphate buffered saline and transferred into a 2 mL screw-top cap. These were heated for 2 hours at 92 °C in a water bath to ensure bacterial killing. Samples were centrifuged at 16000 rpm and pellets washed in 1 mL Milli-Q water 3 times. The resulting pellet was resuspended in 1 mL methanol and transferred into a glass tube. 500 μL chloroform was added to generate 2:1 (v/v) methanol: chloroform solution. This underwent mixing overnight on an automated rotator at room temperature. Next day, samples were centrifuged at 100 g for 10 minutes, and supernatants transferred into new glass vials and stored at room temperature. The pellets were resuspended in 1 mL methanol: 1 mL chloroform, and mixed for 5 hours on the automated rotator and room temperature. Samples were then centrifuged for 5 minutes at 100 g and supernatant was taken and combined with the previous supernatant. These pooled supernatant samples were dried under nitrogen stream then resuspended in 100 μL of dichloromethane and vortexed vigorously. 10 μL per sample was spotted onto silica gel 60 F_254_ TLC plates (Sigma-Aldrich) using a glass syringe. PDIM standard dissolved in dichloromethane was ran as a control. Purified PDIM was provided by American Type Culture Collection, (ATCC), Virginia USA. Plates were ran twice in petroleumether: diethylether (v/v) 9:1. After air drying, plates were stained with 5% phosphomolybdic acid in ethanol (solution at 4 °C) and developed by drying with a heat gun. The TLC was performed twice.

### Volcano Plots

For volcano plots for visualising metabolomic and lipidomic datasets, peaks were automatically integrated on Progenesis and the values normalised to total ion abundance. A filter of co-efficient of variation (CV) < 30 was applied in Progenesis, and only the features meeting this filter were exported to Microsoft Excel. Mean abundance was then calculated across the replicates within each strain/ condition. These mean values were used to calculate fold changes, which are plotted as the log_2_ transformation on the x-axis. To facilitate fold-change calculations, when a feature was below the limit of detection, a missing value of 1000 was assigned.

Student’s t-test was used to compare means between groups and were calculated using the individual values across all replicates within each group. The resulting p-values were then adjusted for multiple comparisons using a false discovery rate approach with a desired FDR of 5%, calculated using GraphPad Prism. The -log_10_ transformation of these values are plotted on the y-axis. All volcano plots were generated using OriginPro. For the negative ion-mode data in the metabolomics dataset in Figure 1a, all features that were significantly altered in the Δ*glpQ1* strain versus the WT were further analysed in Freestyle 1.7 SP2, including the manual integration of the AUC and these values were used, after normalisation to total ion abundance, to generate the volcano plot.

For specific later analyses, following confident feature annotation based on accurate mass, retention time and C.I.D. data, species were no longer excluded if they failed the CV < 30 filter. Therefore more species may have been included in these analyses than appear in the corresponding volcano plot. 5 or more replicate cultures were analysed per strain/condition.

## End Notes

### Author Contributions

R.M.G. devised the experiments, conducted the experiments and wrote the manuscript. L.P.S.d.C devised the project, devised the experiments and edited the manuscript. D.M.H. and A.A. generated the mutant strains of Mtb. M.S.d.S., A.G-G. and J.M. performed the LCMS. J.L. and R.E.L performed the chemical synthesis. A.R. and M.G.G. carried out the mouse experiments under licence.

### Additional Information

Supplementary Information is available for this paper

Correspondance and requests for materials should be addressed to Luiz Pedro Carvalho

Reprints and permissions information is available at www.nature.com/reprints

## Extended Data Legends

**Extended Data Figure 1. Envelope lipids of Mtb.** a: Model of the envelope of Mtb. The envelope comprises of two lipid bilayers, the plasma membrane (PM) of phospholipids and acylated phosphatidylinositol mannosides (AcPIMs) as the inner membrane, and the mycobacterial outer membrane (MOM) made up of longer chain, highly hydrophobic mycolic acids along with other complex lipids, many of which unique to mycobacteria. Between the two is a cell-wall structure of arabinan, galactomannan and peptidoglycan. Long lipoglycans lipomannan (LM) and lipoarabinomannan (LAM) extend from a phosphatidylinositol membrane anchor in the PM out into the periplasm. Finally, a lipid-poor, polysaccharide-rich capsule exists beyond the outer lipid membrane. Ac_1/2_PIM_2/6_: mono/di-acylated phosphatidylinositol di/hexa-mannoside. AGP: arabinogalactan-peptidoglycan. b: The phospholipid recycling pathway in Mtb. The four classes of phospholipid present in the PM undergo hydrolysis of their fatty acyls catalysed by phospholipases to produce their corresponding polar head groups, or lipid-heads. These are hydrolysed by glycerophosphodiesterase activity into the common product glycerol-3 phosphate, and an alcohol. Glycerol-3 phosphate can be further metabolised by glycerol-3 phosphate phosphatase activity, such as by Rv1692, to glycerol and phosphate, or can be re-acylated to phosphatidic acid (PA) and then activated to cytidine diphosphate-diacylglycerol (CDP-DAG) committing to phospholipid synthesis via subsequent class specific reactions. This pathway enables remodelling of the PM by adjustment in the relative amounts of each phospholipid present. PE: phosphatidylethanolamine, PG: phosphatidylglycerol, PI: phosphatidylinositol, CL: cardiolipin. CdsA: phosphatidate cytidylyltransferase. c: Comparison of a typical PM of a Gram-negative bacteria to that of Mtb. Left-bilayer formed almost exclusively of PE and PG. Right, mycobacterial PM: constructed of four conventional phospholipids PE, PG, PI and CL, with AcPIM_2_ and AcPIM_6_ further major components. AcPIMs are unique to actinomycetes. The membrane may be asymmetric, with the inner layer composed predominantly of AcPIM_2_. These bulky lipids reduce the permeability of the membrane. Trehalose monomycolate (TMM) may also be present in the membrane, and is thought to act as a shuttle for mycolic acids across the PM out into the MOM.

**Extended Data Figure 2. Further polar metabolites altered by *glpQ*1 deletion.** a: Volcano plot showing the metabolome of Mtb plotted as the fold change in the mean abundance of each feature in Δ*glpQ*1 strain/parent strain, showing positive ion-mode. Means were calculated across replicate cultures: Δ*glpQ*1 = 6 replicates, parent = 8 replicates. This is the corresponding positive mode dataset do the negative mode data plotted in Figure 1a. 1064 features are plotted, 67 of which of significantly altered in abundance. The canonical lipid-heads of Mtb are highlighted in red. The feature in green is glycerophosphocholine (M+H)^+^ *see Figure 2*. b: Plots of the abundance of each of the four canonical lipid-heads of Mtb in extracts from each strain as labelled, plotted as fold changes with the abundance in the WT set to 1. Data is from the amide column chromatography method. For bis(GroP)Gro the abundance in the WT was below the level of detection, so an arbitrary value of 1000 was assigned to allow fold change to be calculated. c: The structure and MS2 spectrum for glycerophosphothreonine, seen to accumulate in the Δ*glpQ*1 strain in negative ion-mode and not before described in mycobacteria. Key product ions are annotated with their corresponding fragment structures and their expected masses.

**Extended Data Figure 3. Lipidome remodelling in response to *glpQ*1 deletion.** a and b Volcano plots showing the lipidome of Mtb plotted as the mean abundance of each ion feature in the Δ*glpQ*1 strain/ in the parent strain. a: positive ion-mode b: negative ion-mode. Total number of features detected is shown, as well as the percentage of features significantly downregulated (blue) and upregulated (orange) in the Δ*glpQ*1 strain. Fold changes are calculated from means across replicate cultures. Representative of three independent experiments. c: Heat map showing mean fold change in abundance of each individual species of phospholipid in the Δ*glpQ*1 strain versus the parent and in the Δ*glpQ*1::*glpQ*1 strain versus the parent, log_10_ scale. Red indicates enrichment versus the parent strain, blue depletion. Phospholipid species are arranged into their classes on the x-axis. d and e Scatter plots showing the summed abundance of the 5 species of mono- and di-acylated PIM_2_s detected in the lipid extracts from each strain. Points represent replicate cultures, horizontal lines the mean across replicates. d. experiment 1, e. experiment 2. f: Structure of an example mono-acylated PIM_2_. Di-acylated PIMs have a fourth acylation at the hydroxyl group highlighted on the inositol residue. The two acylations of the glycerol do not contribute to the mono- and di-acylation nomenclature, as these form part of the PI anchor and are present in all PIMs.

**Extended Data Figure 4.** Repeat glycerophosphocholine sole phosphate source experiment. Growth profiles of the strains as labelled in Pi-replete (25 mM) and Pi-free (0 mM) media. At day 11 bacteria were transferred from Pi-free media into fresh Pi-replete media or into GroPCho media (0 mM Pi, 25 mM glycerophosphocholine). Duplicate cultures were performed per strain. This is the repeat experiment of the one shown in Figure 2d.

**Extended Data Figure 5. Mouse infection studies.** a. Results of low-dose infection of C57BL/6 mice. Plots of colony forming units per mouse lung (log_10_ scale) for mice infected with either WT Mtb (black), the Δ*glpQ*1 strain (red). 5 mice were sacrificed per timepoint per strain. Points represent the means, error bars the standard error of the mean. Left plot: experiment 1. Right plot: repeat experiment. b: Thin layer chromatography of PDIM extracts from various strains of H37Rv generated by our laboratory. WT is the parent/WT strain used in all the experiments in this study. 1. is the Δ*glpQ*1 strain, and 5. is the Δ*glpQ*1::*glpQ*1 strain. 2, 3, 4, and 6 are strains of H37Rv for which no data appears in this study. The rightmost lane (+) contains purified PDIM standard obtained from b.e.i. resources. Note that strains 1 - 3 and 5 - 6 are all progeny of the parent WT strain used in this study. Strain 4 is a further strain generated from a H37Rv stock several years previously.

**Extended Data Figure 6. Further features of the phosphate starvation lipidome.** a: Total Ion Chromatograms for lipid extracts from WT Mtb cultured in Pi-free media (top) and Pi-replete media (25 mM) (bottom), positive ion-mode. Chromatograms are from single cultures but are highly representative of all replicates across two independent experiments. b: volcano plot showing the lipidome of Mtb plotted as the mean abundance of each ion feature in the WT grown in Pi-free media/ mean abundance in the WT grown in Pi-replete (25 mM) media, negative ion-mode. This is the corresponding negative mode data to the positive ion-mode data plotted in Figure 3a. Representative of 2 independent experiments. PIMs are highlighted in red-purple (horizontal split) and MPMs in red-purple (vertical split). The percentage of features down- and up-regulated are labelled. c: Table showing the mean fold change in abundance of all 70 phospholipids detected, summed by their phospholipid class. For the conventional phospholipids, the area under the curves were summed from positive ion-mode, for PIMs and MPMs negative ion-mode was used as these lipids were not quantifiable in positive ion-mode. Data is from the first of two independent experiments. d. Scatter plots showing the total summed abundance of the 5 species of mono- and di-acylated PIM2 detected in the lipid extracts of the WT grown in Pi-replete media and in Pi-free media, as indicated. Individual points represent individual replicate cultures, horizontal lines the means across replicates. Left plot: experiment 1. Right Plot: repeat experiment. e. Chemical structure of an example MPM (β-D-mannosyl phosphomycoketide). Species vary by the length of the saturated oligoisoprenoid chain, which contains 5 methyl branches. f: Scatter plot showing the total summed abundance of the 4 species of MPM detected in the lipid extracts of the WT grown in Pi-replete media and in Pi-free media, as indicated. Individual points represent individual replicate cultures, horizontal lines the means across replicates. g: EIC overlays for an example of each class of phosphorus-free replacement lipid, with the EICs for each replicate culture of the WT grown in Pi-free media (top) and Pi-replete (25mM) media (bottom) overlain. Scale bars show the intensity of detection in positive ion-mode in counts (arbitrary units). For each plot, the Pi-replete chromatogram is drawn on the same scale as the Pi-free chromatogram.

**Extended Data Figure 7.** Scheme for the chemical synthesis of GlcA-DAG (16:0/16:0). Reagents and reaction conditions are labelled. See *methods* for full details.

