## Extended Data Figures for "*Mycobacterium tuberculosis* overcomes phosphate starvation by extensively remodelling its lipidome with phosphorus-free lipids"

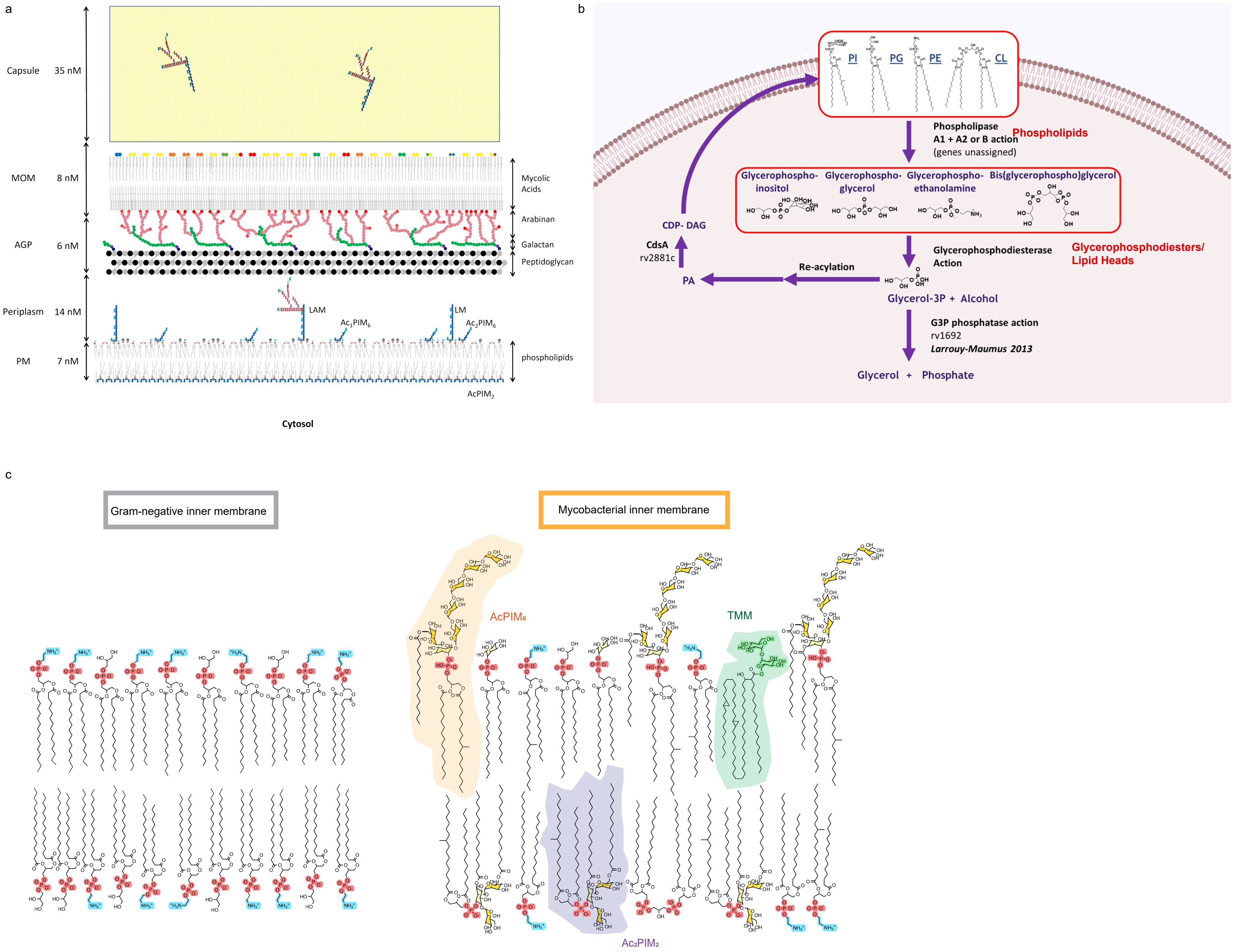
a
Capsule 35 nM
MOM 8 nM
AGP 6 nM
Periplasm 14 nM
PM 7 nM
Mycolic Acids
Arabinan
Galactan
Peptidoglycan
LAM
LM
Ac1PIM6
Ac2PIM6
phospholipids
AcPIM2
Cytosol
b
PI
PG
PE
CL
Phospholipase A1 + A2 or B action (genes unassigned)
Phospholipids
Glycerophospho-inositol
Glycerophospho-glycerol
Glycerophospho-ethanolamine
Bis(glycerophospho)glycerol
CDP- DAG
CdsA rv2881c
Glycerophosphodiesterase Action
Glycerophosphodiesters/ Lipid Heads
Re-acylation
PA
Glycerol-3P + Alcohol
G3P phosphatase action
rv1692
Larrouy-Maumus 2013
Glycerol + Phosphate
c
Gram-negative inner membrane
Mycobacterial inner membrane
AcPIM6
TMM
Ac2PIM2

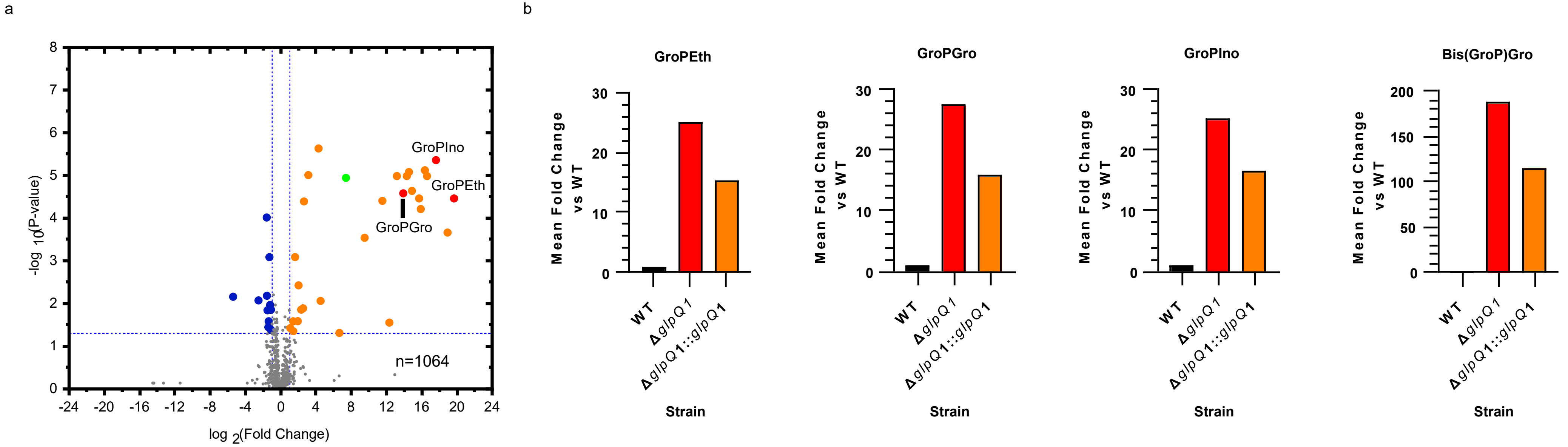


c

**Glycerophosphothreonine, GroPThr**
**$C_7H_{15}NO_8P$-**
**m/z $(M-H)^-$ 272.0541 (calculated)**
**272.0539 (observed)**

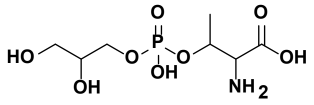


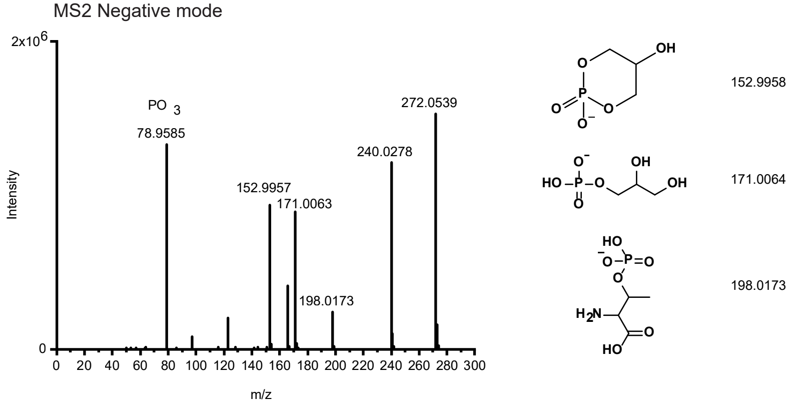

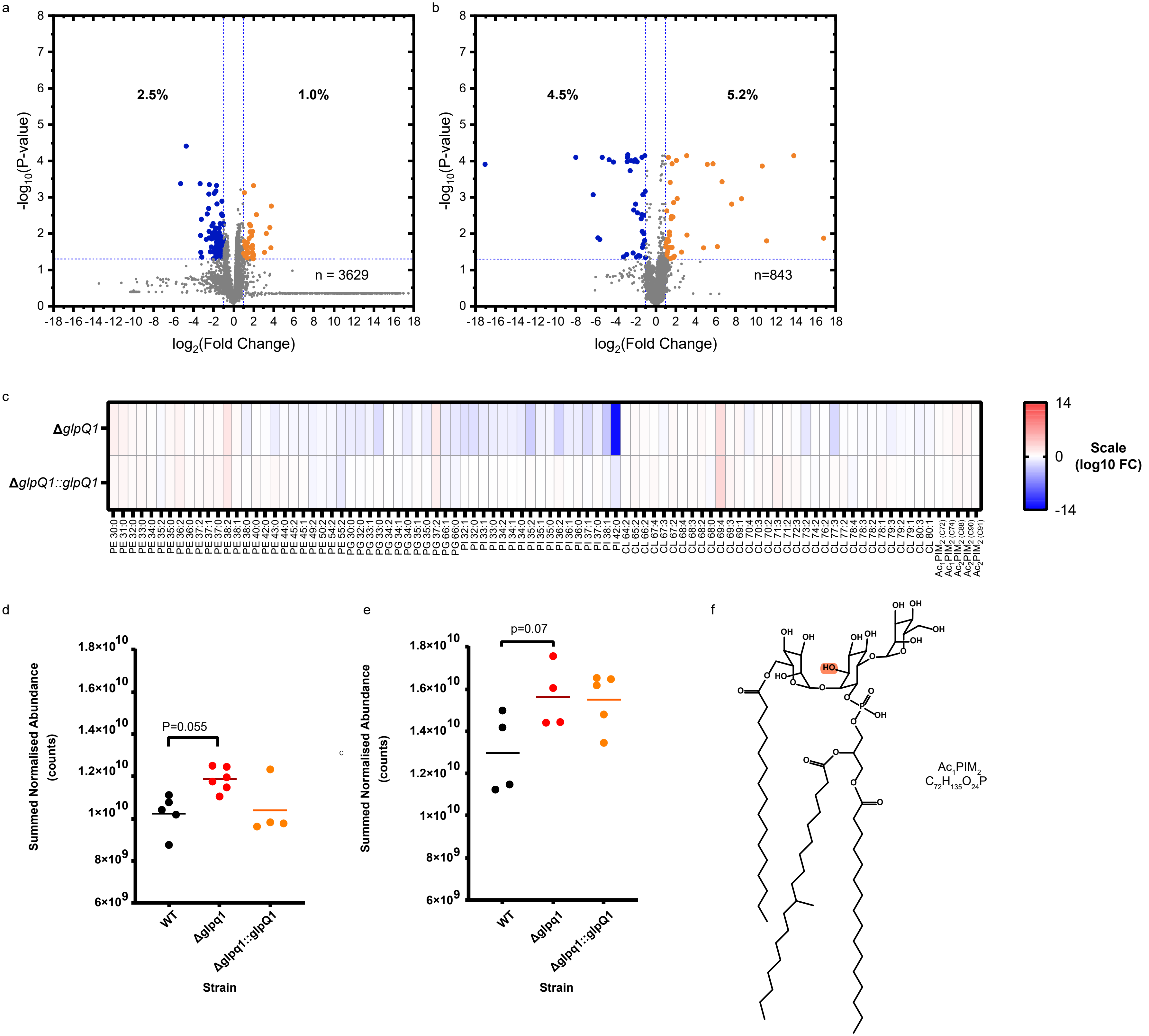
a
2.5%
1.0%
-log10(P-value)
log2(Fold Change)
n = 3629
b
4.5%
5.2%
-log10(P-value)
log2(Fold Change)
n=843
c
ΔglpQ1
ΔglpQ1::glpQ1
PE 30:0
PE 31:0
PE 32:0
PE 33:0
PE 34:0
PE 35:2
PE 35:0
PE 36:2
PE 36:0
PE 37:2
PE 37:1
PE 37:0
PE 38:2
PE 38:1
PE 38:0
PE 40:0
PE 42:0
PE 43:0
PE 44:0
PE 45:2
PE 45:1
PE 49:2
PE 50:2
PE 54:2
PE 55:2
PG 30:0
PG 32:0
PG 33:1
PG 33:0
PG 34:2
PG 34:1
PG 34:0
PG 35:1
PG 35:0
PG 37:2
PG 66:1
PG 66:0
PI 32:1
PI 32:0
PI 33:1
PI 33:0
PI 34:2
PI 34:1
PI 34:0
PI 35:2
PI 35:1
PI 35:0
PI 36:2
PI 36:1
PI 36:0
PI 37:1
PI 37:0
PI 38:1
PI 42:0
CL 64:2
CL 65:2
CL 66:2
CL 67:4
CL 67:3
CL 67:2
CL 68:4
CL 68:3
CL 68:2
CL 68:0
CL 69:4
CL 69:3
CL 69:1
CL 70:4
CL 70:3
CL 70:2
CL 71:3
CL 71:2
CL 72:3
CL 73:2
CL 74:2
CL 76:2
CL 77:3
CL 77:2
CL 78:4
CL 78:3
CL 78:2
CL 78:1
CL 79:3
CL 79:2
CL 79:1
CL 80:3
CL 80:1
Ac1PIM2 (C72)
Ac1PIM2 (C74)
Ac2PIM2 (C88)
Ac2PIM2 (C90)
Ac2PIM2 (C91)
14
0
-14
Scale
(log10 FC)
d
Summed Normalised Abundance
(counts)
P=0.055
WT
Δglpq1
Δglpq1::glpQ1
Strain
e
Summed Normalised Abundance
(counts)
p=0.07
WT
Δglpq1
Δglpq1::glpQ1
Strain
f
Ac1PIM2
C72H135O24P

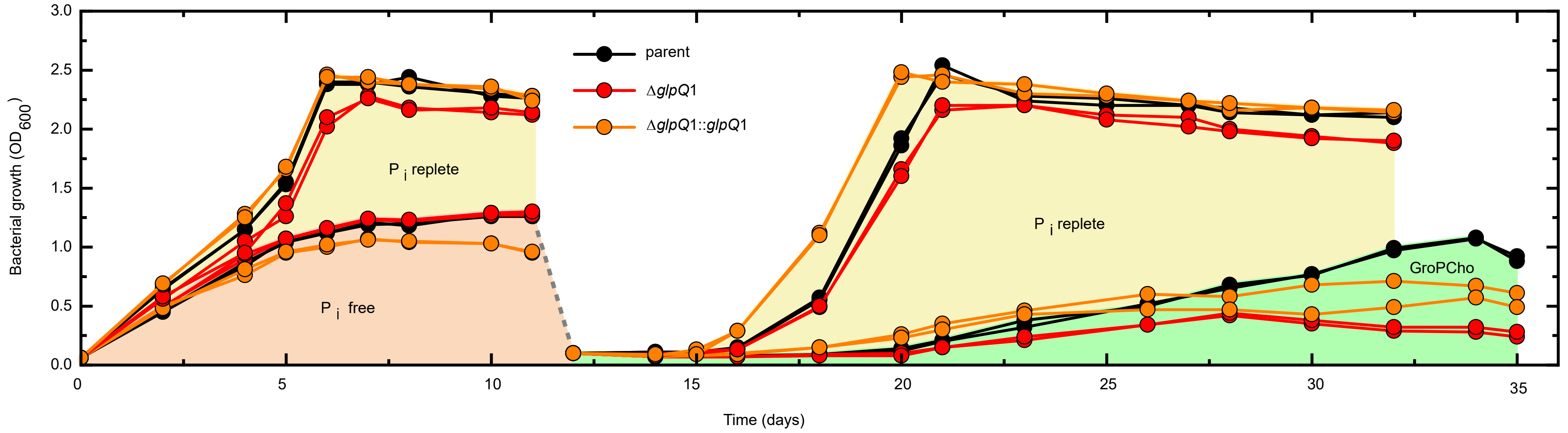

parent
ΔglpQ1
ΔglpQ1::glpQ1
P i replete
P i free
P i replete
GroPCho
Bacterial growth (OD600)
Time (days)
3.0
2.5
2.0
1.5
1.0
0.5
0.0
0
5
10
15
20
25
30
35

a

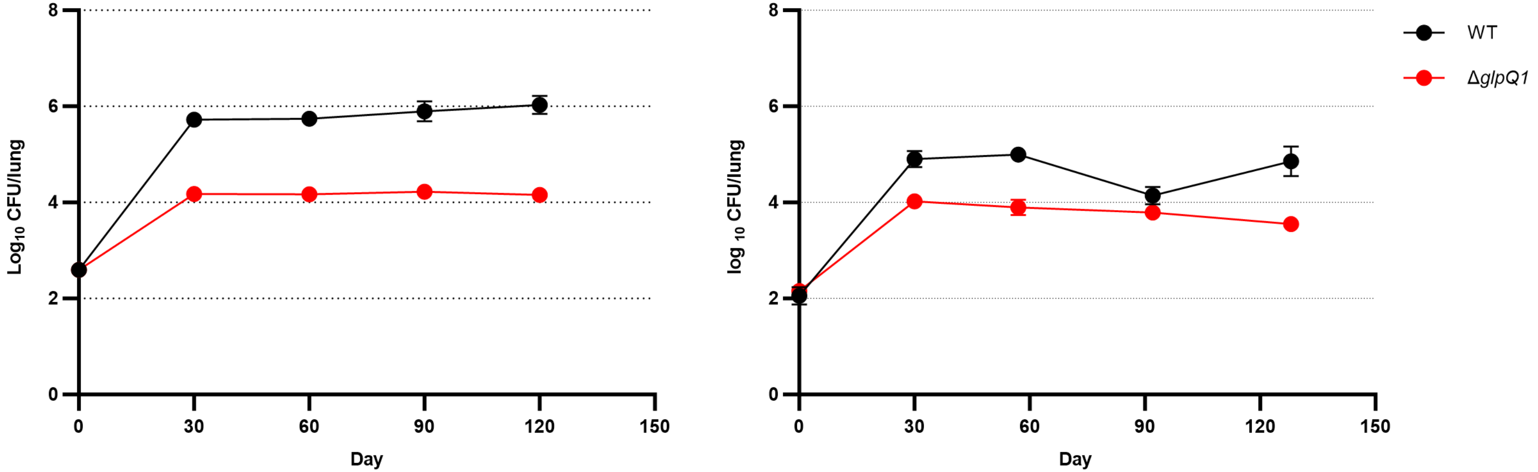

Log10 CFU/lung
log 10 CFU/lung
Day
Day
8
6
4
2
0
0
30
60
90
120
150
WT
ΔglpQ1


b

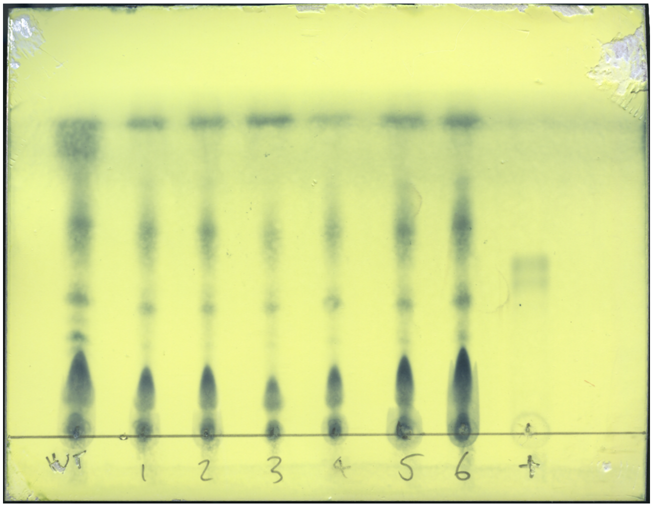

WT
1
2
3
4
5
6

a

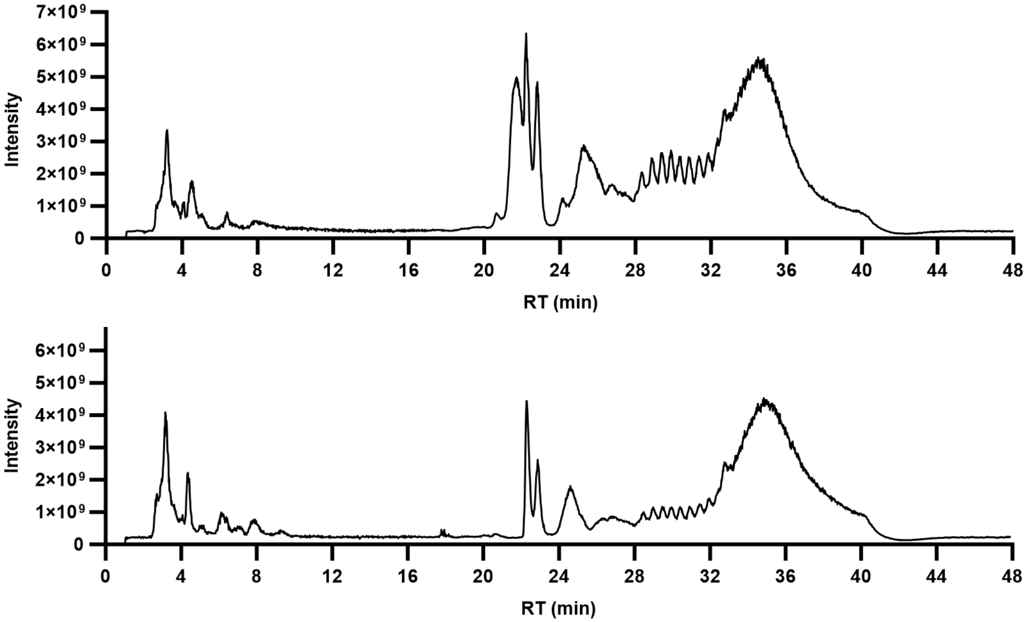


b

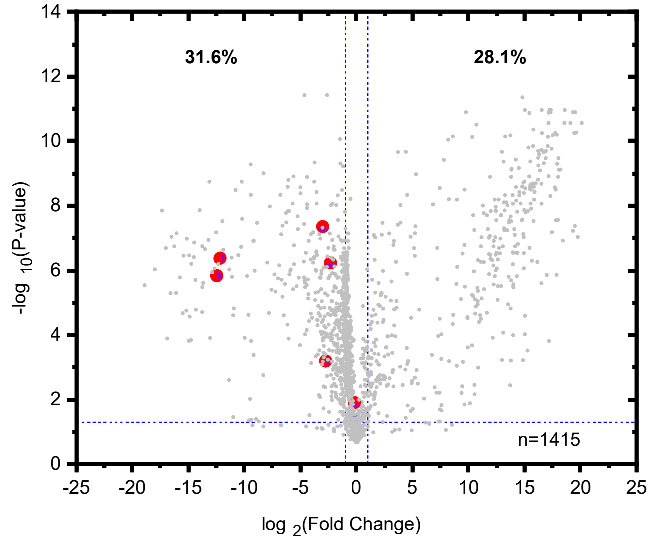


c

| Class | no. of lipids | Total area under curve (average across replicates) | | Fold change | T-test |
|---|---|---|---|---|---|
| | | Pi free | Pi free | Pi free / Pi replete | p value |
| PEs | 17 | 9.49 x$10^{8}$ | 3.88 x$10^{10}$ | 0.024 | 7.2 x$10^{-11}$ |
| PGs | 16 | 9.12 x$10^{8}$ | 4.28x$10^{8}$ | 2.134 | 3.1x$10^{-7}$ |
| PIs | 8 | 1.92 x$10^{9}$ | 8.84x$10^{9}$ | 0.217 | 9.4 x$10^{-4}$ |
| CLs | 20 | 1.46 x$10^{10}$ | 1.95x$10^{10}$ | 0.750 | 5.7 x$10^{-5}$ |
| All conventional PLs | 61 | 1.84x$10^{10}$ | 6.75x$10^{10}$ | 0.272 | 1.1x$10^{-10}$ |
| PIM | 5 | 3.17 x$10^{9}$ | 4.75 x$10^{9}$ | 0.669 | 1.0 x$10^{-4}$ |
| MPM | 4 | 2.37 x$10^{6}$ | 4.23x$10^{7}$ | 0.056 | 4.8 x$10^{-8}$ |
| All PLs | 70 | 2.16 x$10^{10}$ | 7.23 x$10^{10}$ | 0.298 | 1.1x$10^{-10}$ |

d

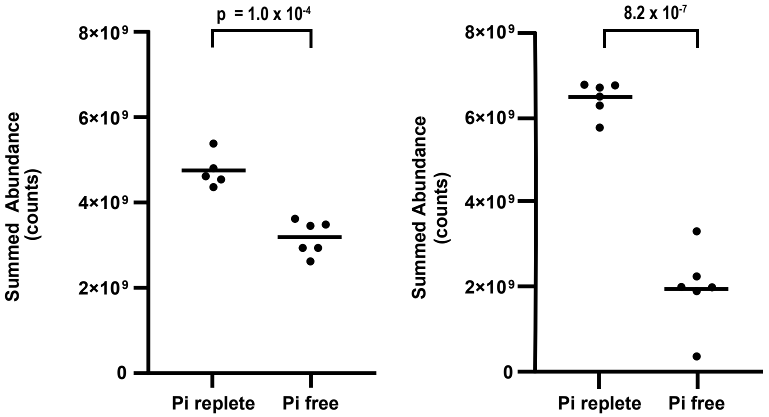


e

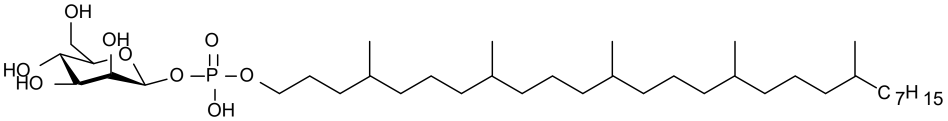


f

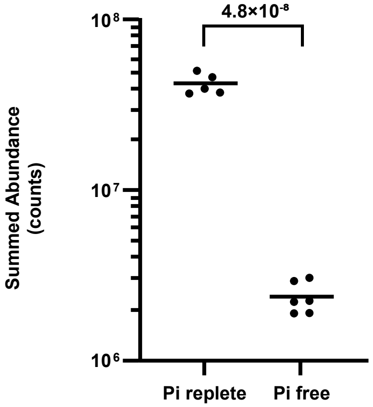


g

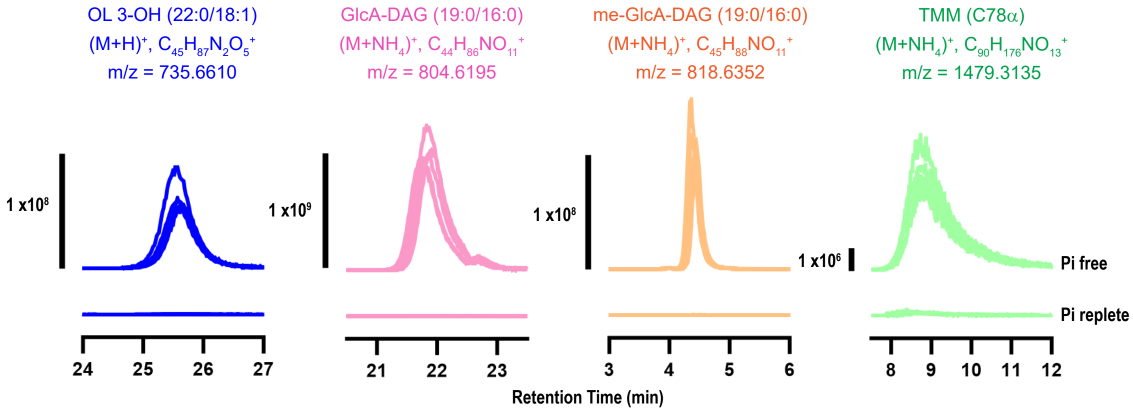

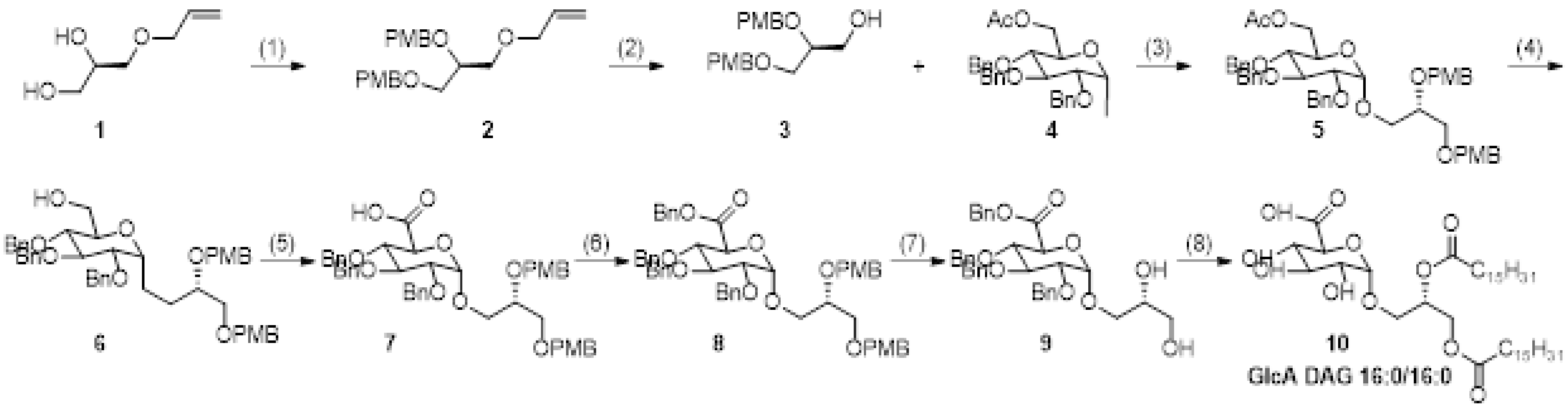


Reagents and conditions: (1) NaH 95%, DMF, p-methoxybenzyl chloride, TBAI, 71%; (2) PdCl2, CuCl, DMF, 44%; (3) TBAI, 2,4,6-tri-tert-butylpyrimidine, CH2Cl2, 60%; (4) NaOMe in MeOH, CH2Cl2, 92%; (5) TEMPO, BAIB, CH2Cl2/H2O, 88%; (6) benzyl alcohol, HBTU, DIPEA, DMAP, CH2Cl2, 34%; (7) CAN, MeCN/H2O, 44%; (8) a. palmitic acid, COMU, DMAP, DMF; b. Pd(OH)2/C, H2, MeOH/THF, 8%.
